# Possible acquisition and molecular evolution of *vpu* genes inferred from comprehensive sequence analysis of human and simian immunodeficiency viruses

**DOI:** 10.1101/2024.04.03.587873

**Authors:** Miu Naruki, Motofumi Saito, Masako Nomaguchi, Akio Kanai

**Affiliations:** Institute for Advanced Biosciences, Keio University, Tsuruoka 997-0017, Japan; Systems Biology Program, Graduate School of Media and Governance, Keio University, Fujisawa 252-0882, Japan; Department of Microbiology, Graduate School of Medicine, Tokushima University, Tokushima 770-8503, Japan; Faculty of Environment and Information Studies, Keio University, Fujisawa 252-0882, Japan

**Keywords:** HIV-1, SIV, Vpu, bioinformatic, phylogenetic tree, molecular evolution

## Abstract

Vpu, an accessory protein of *Human immunodeficiency virus 1* (HIV-1), plays a crucial role in viral particle production and release, contributing to HIV virulence and its associated pandemic. Despite its significance, the acquisition and evolution of the *vpu* gene remain poorly understood. In this study, we conducted a comprehensive computational analysis of approximately 38,000 sequences from *Simian immunodeficiency viruses* (SIV) and HIV. From these, we obtained 141 representative Vpu protein sequences, while preserving their diversity by removing redundant sequences. We then created a phylogenetic tree, which revealed that SIV and HIV strains can be classified into four major types based on the Vpu proteins they express: Vpu-type 1, the putative ancestral type, which includes SIVs from Dent’s mona monkey, mona monkey, and moustached guenon; Vpu-type 2, which includes SIVgor from gorillas and HIV-1 group O; Vpu-type 3, which includes SIVcpz from chimpanzees; and Vpu-type 4, which includes HIV-1 group M, HIV-1 pandemic strain (subtype 4a), and HIV-1 group N (subtype 4b). Vpu-types 1 and 2 show variations in *vpu* length, whereas Vpu-types 3 and 4 show less variability. Notably, the gene encoding Vpu-type 1 varies in the length of its overlap with the *env* gene. To clarify the evolutionary relationship between Vpu-type 1 SIVs and SIVs lacking Vpu, we constructed a phylogenetic tree based on the nucleotide sequence between the *pol* and *env* genes, using 401 sequences derived from HIV-1, HIV-2, and SIVs. Vpu-type 1 (SIVden and SIVmon), SIVasc (red-tailed monkey), and SIVsyk (Sykes’ monkeys) clustered closely on the phylogenetic tree. Considering the observed similarities between the *vpu, vpr* and *env* genes in SIVdeb and SIVsyk, we speculated that the *vpu* gene originated within the SIV genome. A phylogenetic tree constructed with 242 Vpu-type 4a sequences from the HIV pandemic strain and 135 sequences of circulating recombinant forms of HIV-1 revealed that the viral subtypes can be further classified into 18 distinct protein subtypes, surpassing the number of previously known subtypes. Collectively, our results suggest an evolutionary model for the *vpu* gene that sheds light on its origin and divergence, and their implications in the context of HIV-1 and SIV evolution.

## 1. Introduction

*Human immunodeficiency virus 1* (HIV-1) is an RNA virus in the genus *Lentivirus* and the family *Retroviridae*. HIV emerged through zoonotic cross-species transmission from *Simian immunodeficiency virus* (SIV) infecting primates in Africa (Kirchhoff, 2009). Currently, over 40 nonhuman primate species carry species-specific SIV infections (Aghokeng et al., 2010). HIV-1 is categorized into four groups: group M (main), group O (outlier), group N (non-M/non-O), and group P, among which group M is the pandemic strain. Studies have shown that HIV-1 groups M and N evolved from SIVcpz found in the chimpanzee (*Pan troglodytes*) whereas groups O and P evolved from SIVgor infecting the gorilla (*Gorilla gorilla*) (Keele et al., 2006; Takehisa et al., 2009; Van Heuverswyn et al., 2006). In contrast, HIV-2 evolved from SIVsmm infecting the sooty mangabey (*Cereocebus atys*) (Hirsch et al., 1989). Ancestral differences between HIV-1 and HIV-2 underpin their genetic differences and the differences in their infectivity rates.

SIVs such as SIVgor originated from SIVcpz, whereas SIVcpz evolved from multiple recombination events among SIVrcm, SIVgsn, SIVmon, and SIVmus, which infect red-capped mangabeys, greater spot-nosed monkey (*Cercopithecus nictitans*), mona monkey (*C. mona*), and moustached guenon (*C. cephus*), respectively. These are commonly categorized as Old World monkeys (Sharp et al., 2005). The genome organization of HIV and SIV includes 10 genes (Supplementary Fig. S1): three core genes (*gag*, *pol*, and *env*), two essential regulatory genes (*tat* and *rev*) and five accessory regulatory genes (*vif*, *vpr*, *nef*, *vpx*, and *vpu*). HIV-1 and several SIVs strain, sharing a common evolutionary lineage, contain the *vpu* gene. In contrast, HIV-2 and certain SIVs lack this gene, instead carrying the *vpx* gene or neither of these genes. To extend our understanding of the genetic changes that have occurred in HIV-1 over time, we investigated the origin and evolution of the *vpu* gene in HIV-1. This may also provide insights into the mechanisms of gene acquisition and adaptation in HIV-1, potentially clarifying why one specific strain has emerged as the predominant variant in the global HIV pandemic.

Vpu (viral protein U) is an 81-amino-acid type 1 integral membrane phosphoprotein that is translated by a Rev-regulated bicisotronic mRNA that also encodes the *env* gene (Cohen et al., 1988; Schwartz et al., 1990; Strebel et al., 1989). Vpu comprises three distinct α-helices: the N-terminal proximal transmembrane domain and two C-terminal domains (Cohen et al., 1988; Federau et al., 1996; Gonzalez, 2015; Schubert et al., 1996). Its functions are to enhance the degradation of CD4^+^ T cells in the endoplasmic reticulum and augment the release of progeny virions from infected cells by antagonizing tetherin (also referred to as BST-2). A host restriction factor enhances the infectivity and pathogenesis of the virus overall (Klimkait et al., 1990; Schubert & Strebel, 1994; Terwilliger et al., 1989; Willey et al., 1992). It is noteworthy that only HIV-1 group M strains contain a *vpu* gene that expresses Vpu with the dual capacity to enhance the degradation CD4^+^ T cells and antagonize tetherin effectively. The N, O, and P groups of HIV-1 do not show this trait (Sauter et al., 2009). The acquisition of this fully functional Vpu in HIV-1 group M is thought to have contributed to the global spread of the pandemic (Volcic et al., 2020). However, the origin of the *vpu* gene and the mechanisms underlying its molecular evolution in HIV-1 are still poorly understood. A previous study suggested that HIV-1 *vpu* originated from a common ancestor of the gene in SIVgsn, SIVmus, and SIVmon, and was subsequently transferred to the SIVcpz genome (Bibollet-Ruche et al., 2004). However, no study has tested this hypothesis.

In this study, we comprehensively examined an extensive dataset of HIV and SIV sequences. A systematic approach was used to clarify the origin of the gene and trace the evolution of HIV-1 *vpu*. We also performed an evolutionary analysis to understand how the *vpu* gene of the HIV pandemic strain diversified over time and evolved within HIV-1 group M. This investigation has led to a more-detailed reclassification of the subtypes based on their Vpu amino acid sequences. In this study, we also considered the variant B strain, known for its increased virulence relative to that of standard subtype B. Overall, the study extends our understanding of the evolution of the *vpu* gene by establishing an evolutionary model. We provide insights into the origin of *vpu*, the possible mechanism of gene acquisition, and the diversification of *vpu* in the pandemic strain of HIV-1.

## 2. Materials and Methods

### 2.1 Dataset

The nucleotide and amino acid sequences of the HIV and SIV strains used in this study were obtained from Los Alamos HIV Sequence Database (https://www.hiv.lanl.gov/, last accessed 22 October 2021), together with details of the sampling region and sampling year. The recently discovered variant B strain (GenBank IDs: MT458931–MT458935 and MW689459–MW689470; MW689465 and MW689466) was extracted from the National Center for Biotechnology Information (NCBI) GenBank (https://www.ncbi.nlm.nih.gov/). In total, 10,891 nucleotide sequences of viral genomes, 38,841 Vpu amino acid sequences, and 19,965 Pol amino acid sequences were extracted (Supplementary Table S1).

### 2.2 Multiple-sequence alignment

The nucleotide and amino acid sequences of the HIV-1 and SIV strains were aligned with MAFFT L-INS-i v7.45 (Katoh & Standley, 2013) using the default parameters. The alignments were visualized with Jalview v2.11 (Waterhouse et al., 2009). The alignment was colored by ClustalX and the percentage identity color scheme was utilized to highlight the percentage abundance of aligned residues and the amino acids conserved across sequences (Waterhouse et al., 2009). The conservation, consensus, and multi-Harmony scores are shown below the alignments. To quantitatively measure conservation, the numbers of amino acids with specific physicochemical properties conserved in each column of the alignment are calculated. This index is based on the Analysis of Multiply Aligned Sequences (AMAS) method (Livingstone & Barton, 1993), where conservation is represented by a numerical score that reflects the degree of similarity between the physicochemical properties of the amino acids in the alignment, followed by the highest identity score, and substitutions to amino acids in the same physicochemical class. The conservation score for each amino acid position ranges from 1 to 11. High conservation on the alignment is marked with ‘*’ (a score of 11 on the default amino acid grouping), whereas partial conservation in all properties is marked with a yellow ‘+’ (a score of 10) (Livingstone & Barton, 1993). The consensus annotation shows the percentage of the modal residue per column. We used ‘+’ to show that the modal value is shared by more than one residue. Multi-Harmony (Brandt et al., 2010) was used to identify significant patterns of subgroup variation among the columns of an alignment and specific residues of each type. The tool requires that the alignment is subdivided into groups containing at least two nonidentical protein sequences, which we grouped according to the viral type. The Sequence Harmony (SH) scores for each subgrouped alignment were determined by applying Shannon’s entropy (Shannon, 1948), which measures the extent of the differences in amino acid composition between groups, thus detecting subtype-specific sites.

### 2.3 Molecular phylogenetic analysis

CD-HIT is a clustering program used to minimize the number of redundant sequences encoding similar proteins (Fu et al., 2012). It allows researchers to create a dataset composed of solely unique representative sequences. The alignment file created in this analysis was used to construct the phylogenetic tree. trimAl version 1.2rev59 (Capella-Gutierrez et al., 2009) was used to trim the gaps in the alignment. Midpoint-rooted and unrooted maximum likelihood trees were constructed using IQ-TREE v.1.6.12 (Minh et al., 2020). Ultrafast bootstrap approximation (UFBoot2) was applied (Hoang et al., 2018), which allows the rapid and unbiased estimation of branch support values, with 1,000 bootstrap replicates. The best-fit model for the analysis was determined with ModelFinder (Kalyaanamoorthy et al., 2017), which was used to optimize the tree’s accuracy. FigTree v.1.44 was used to visualize and observe the tree (http://tree.bio.ed.ac.uk/software/figtree/).

### 2.4 Pictures used for the evolutionary model

In our evolutionary model, we incorporated images of multiple monkeys’ faces, after cropping them from their original sizes. The photographs were obtained from Wikipedia Commons, and were originally taken by Michael Gäbler, Alena Houšková, Thomas Springer, Peggy Motsch, Laetitia C, Paul Harrison, Wookiemedia, Aaron Logan, and Jack Hynes, distributed under CC BY 3.0, CC0 1.0, CC BY-SA 3.0, CC BY-SA 4.0, and CC BY 2.5 licenses.

## 3. Results and Discussion

### 3.1 Classification of HIV and SIV Vpu proteins

To comprehensively understand the acquisition and evolution of the Vpu proteins in the genus *Lentivirus*, we collected all available sequences from the Los Alamos HIV Sequence Database. Our dataset included a total of 38,841 sequences, containing 38,792 HIV-1 and 49 SIV Vpu sequences. We utilized CD-HIT to cluster sequences with 90% sequence similarity, resulting in 141 HIV-1 and 16 SIV sequences after removing redundancy. This process ensured that only unique representatives of strains with numerous similar sequences were retained (see Supplementary Table S1). To construct a phylogenetic tree from the Vpu amino acid sequences, we classified the Vpu-containing viruses into four distinct groups: Vpu-types 1, 2, 3, and 4 (Fig. 1). The groups were categorized using three primary criteria of the Vpu proteins. First, we examined the phylogenetic tree structure and identified distinct clades within the Vpu protein sequences. Second, we considered the motifs within the amino acid sequences that are associated with specific functional properties (described in detail below). Third, we assessed the differences in the viral host preferences shown by the various Vpu types. Using these three criteria, we comprehensively classified and characterized the different types of Vpu proteins. Vpu-type 1 strains include SIVden, SIVgsn, SIVmon, and SIVmus, which infect Dent’s mona monkey, the greater spot-nosed monkey, the mona monkey, and the moustached guenon, respectively. Vpu-type 2 strains include SIVgor and HIV-1 groups O and P, which infect gorillas and humans, respectively. The Vpu-type 3 strain is SIVcpz, which infects chimpanzees. Vpu-type 4 strains include HIV-1 groups M and N, both of which infect humans. Although the classification of Vpu-types 1–4 does not necessarily indicate the chronological evolutionary order of the viruses, our results provide a clear framework for understanding the evolutionary relationships among the Vpu-containing viruses. We distinguished Vpu-type 3 and Vpu-type 4 based on the difference in the viral host (chimpanzee and human, respectively) and separated the Vpu-types based on their respective origins. The phylogenetic tree also suggested that chimpanzee Vpu-type 3 gave rise to human Vpu-type 4. The results of our classifications are largely consistent with previous studies, which demonstrated that HIV-1 group O originated from SIVgor and HIV-1 groups M and N originated from SIVcpz (Gao et al., 1999). SIVcpz is believed to have originated from multiple recombination events between various SIV strains, including SIVgsn, SIVmus, and SIVmon (Bailes et al., 2003).

**Figure 1.**
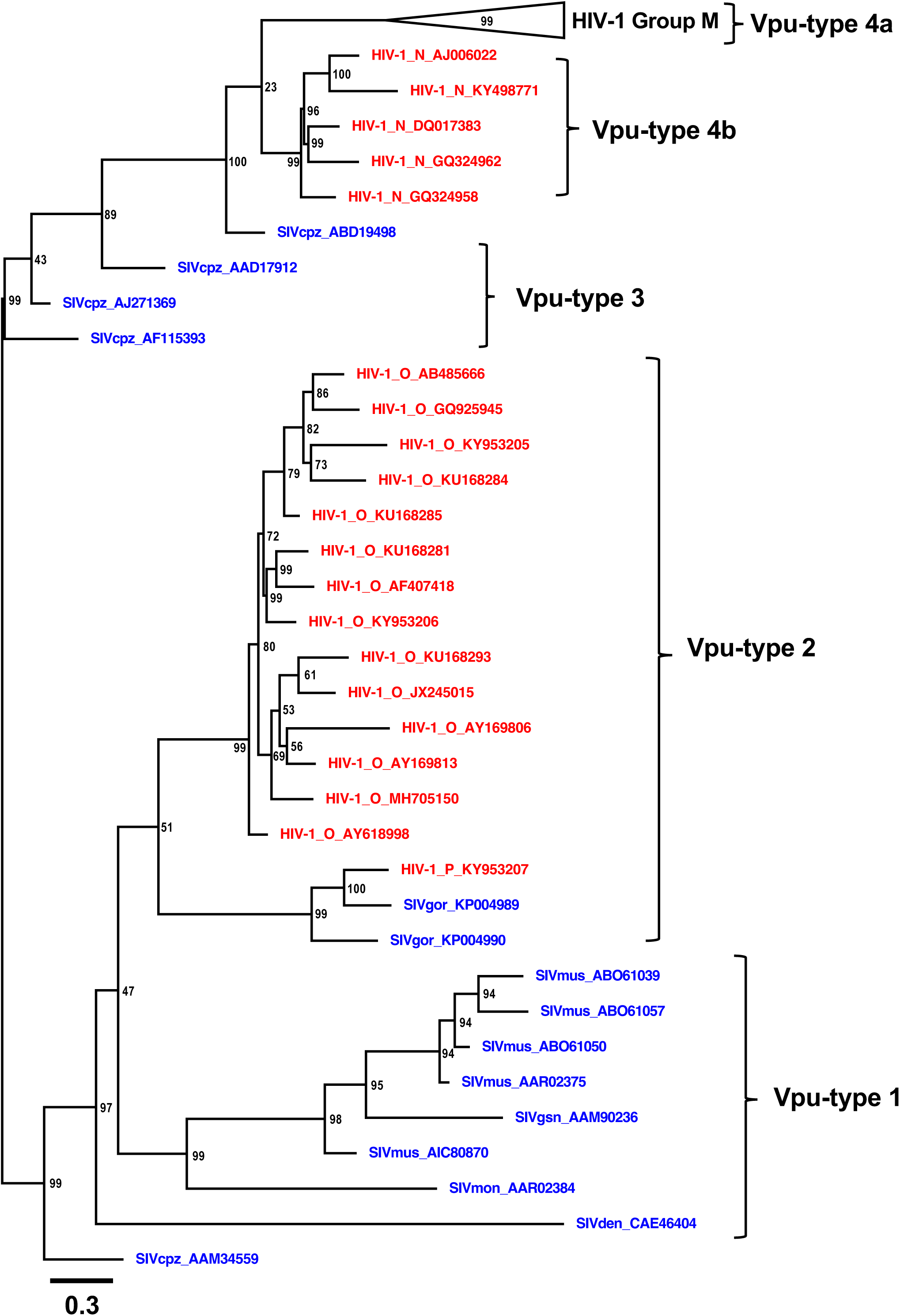
Rooted phylogenetic tree of Vpu proteins from representative HIV-1 and SIV strains. A midpoint-rooted phylogenetic tree of Vpu protein sequences (n = 141; see Supplementary Table S1) was constructed (1,000 bootstrap replicates). The annotations are presented in the order: viral strain, viral group, and GenBank protein ID; and are colored according to the viral host: human (red), apes and monkeys (blue). HIV-1 group M sequences were collapsed and are shown as the black triangle on the tree. Five types of Vpu proteins (Vpu-types 1, 2, 3, 4a, and 4b) were classified based on the clustering of the evolutionary lineages. The scale bar below the tree indicates 0.3 (30%) amino acid substitutions per site, and the bootstrap values (1/100), indicating the estimated posterior probabilities, are shown at each node.

To analyze the differences among the Vpu-types at the amino acid level, we generated a multiple-amino-acid-sequence alignment for each Vpu-type (Fig. 2 and Supplementary Fig. S2). Fig. 2A shows the structure of Vpu, which includes a transmembrane and a cytoplasmic domain, which consists of two α-helices. We established the 81 amino acid positions as a standardized reference for comparison and analysis in this study. This corresponds to the average length of HIV-1 group M Vpu and provided a consistent baseline for our study. In this analysis, we further categorized Vpu-type 4 into two distinct subtypes based on their characteristics: 4a, which includes HIV-1 group M, and 4b, which includes HIV-1 group N. Vpu-type 4b lacks the ability to downregulate CD4^+^ T cells (Sauter et al., 2009; Sauter et al., 2012). Supplementary Fig. S2 shows the overall amino acid alignment of 50 sequences, after randomly selecting 10 sequences from each Vpu-type. The alignment process introduced gaps between within the sequences, resulting in variations in the actual positions of the amino acid residues. Therefore, we refer to the amino acid position indicated in the figure as ‘position’, which takes into account the gaps, and use the term ‘residue’ to refer to the specific amino acid residue without the gaps. Multi-Harmony analysis was conducted to detect and analyze the variations within a set of sequences. For example, we observed that at position 29 (residue 23), the conserved amino acids differed among the Vpu-types. Vpu-types 2, 3, and 4 showed a high degree of conservation (100%), with tryptophan (W) at this position. However, variability was observed in Vpu-type 1 because 20% of the sequences contained tryptophan (W), 40% contained phenylalanine (F), and 40% contained leucine (L). Similarly, at position 30 (residue 24), in Vpu-type 3, glycine (G) was conserved in 70% of sequences, whereas threonine (T) was conserved in 100% of Vpu-type 4a sequences. However, 80% of Vpu-type 4b sequences contained valine (V). At position 59 (residue 42), the following conservation patterns were observed: valine (V) in 70% of type 1 sequences, arginine (R) in 100% of Vpu-type 2 sequences, leucine (L) in 100% of Vpu-types 3 and 4b sequences, and isoleucine (I) in 90% of the sequences between the same position. These findings provide clear evidence of the conservation of specific amino acids within each Vpu-type, highlighting the distinct nature of each type. The alignment also revealed that the C-terminal regions of the sequences are less conserved than the transmembrane domains.

**Figure 2.**
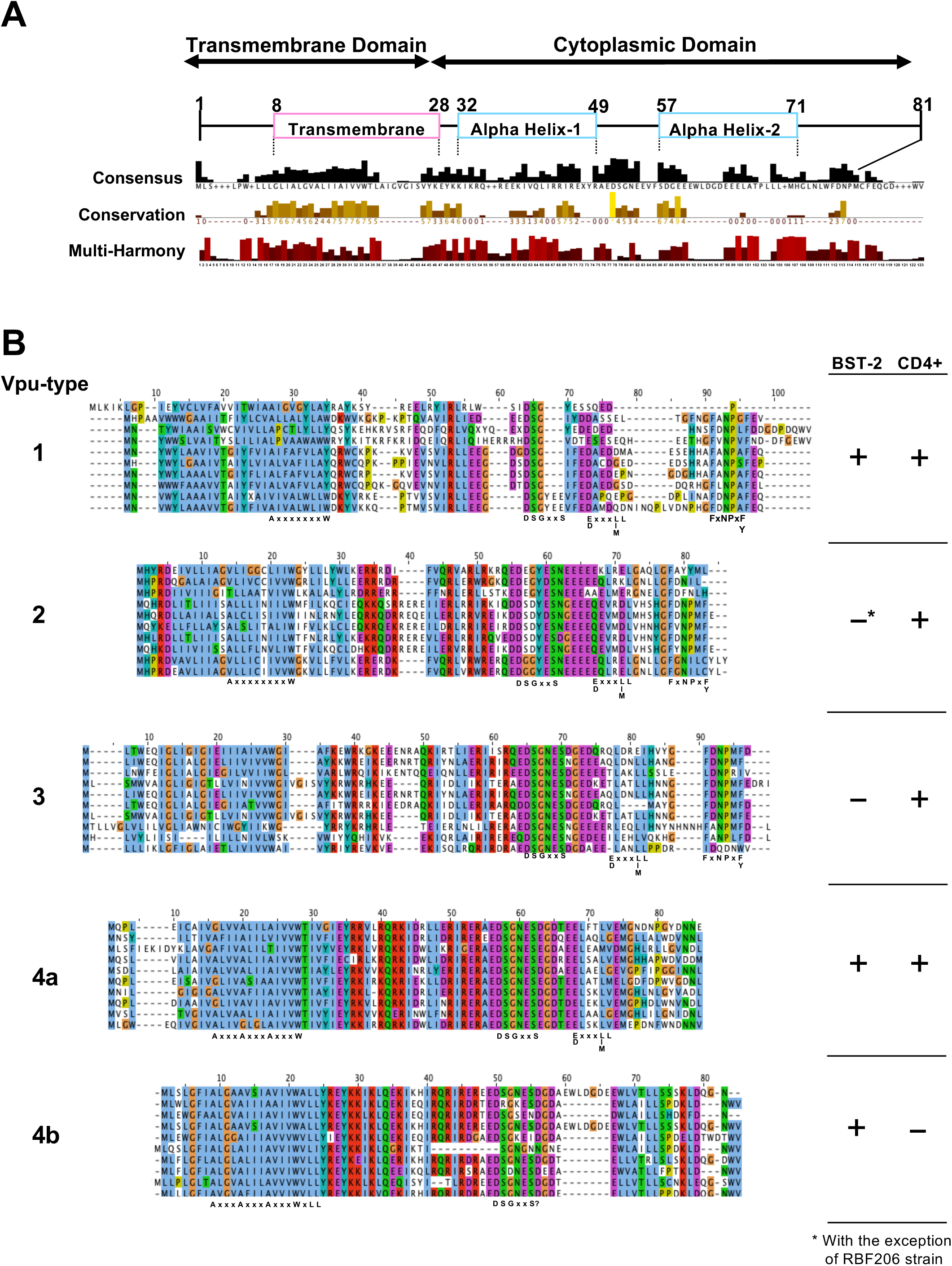
Multiple-amino-acid-sequence alignment of the Vpu proteins. (A) Protein structure of Vpu. The transmembrane domain and cytoplasmic domain regions in the Vpu protein are marked. The consensus sequence displays the conserved regions. Nonconserved residues are marked with ‘+’. The conservation histogram shows the conservation of residues according to their physicochemical properties. The multi-Harmony histogram shows the subtype-specific residues. (B) A multiple-sequence alignment (n = 50) of each type of Vpu protein, colored according to the ClustalX color scheme. The name of the virus containing the corresponding Vpu protein is also given. Important functional motifs and residues are indicated below the alignment. The table on the right shows the ability (+) or inability (−) of each type to antagonize tetherin (BST-2) or degrade CD4^+^ T cells. *Strain RBF206 is the only strain in group O known to be active against tetherin (Mack et al., 2017).

In Fig. 2B, we examined the Vpu-types individually to investigate the Vpu-type-specific differences in detail and to consider the presence or absence of specific functions associated with each Vpu type. These functions included the ability to antagonize tetherin and to downregulate CD4^+^ T cells. The transmembrane domain contains motifs involved in tetherin antagonism, whereas the cytoplasmic domain contains motifs related to the downregulation of CD4^+^ T cells. It has been established that the AxxxAxxxAxxxW, AxxxAxxxAxLL, and AxxxxxxxW residues in the transmembrane domain play a crucial role in the anti-tetherin activity of HIV-1 groups M (Vpu-type 4a) and N (Vpu-type 4b) and SIVgsn (Vpu-type 1), respectively (Douglas et al., 2013; Sauter et al., 2012; Yao et al., 2022). Interestingly, the DSGxxS motif, which is crucial for the proteasomal degradation of CD4^+^ T cells, is partly conserved in Vpu-type 1 because it contains only one of the double serine residues required for this function. Similarly, in Vpu-type 2, the serine (S) residue at position 57 (residue 53) is conserved in 50% of sequences, whereas the remaining sequences contain either a glycine (G) residue (20%) or a glutamic acid (E) residue (30%). However, the serine residue at position 61 (residue 57) is fully conserved across all sequences. In contrast, both S residues located at positions 72 and 79 in Supplementary Fig. S2 are completely conserved throughout Vpu-types 3 and 4. Furthermore, the FxNPxF/Y motif located in the cytoplasmic domain, whose function remains unknown, is commonly found in SIVcpz and HIV-1 group O. Interestingly, we also observed its presence in almost every Vpu-type 1 sequence (Kluge et al., 2013). This phenylalanine (F) is highly conserved throughout Vpu-types 1, 2 and 3, but this residue and motif are notably absent from Vpu-type 4.

Despite the absence of crucial functions in Vpu-type 1, it retains the ability to antagonize both CD4^+^ T cells and tetherin, indicating that the variation in the essential motifs across the different Vpu types does not necessarily affect their functionalities. We also noted variations in the sequence lengths of the different Vpu types. Therefore, we quantitatively analyzed the differences among the Vpu types in terms of the length of the *vpu* gene, the length of overlap between *vpu* and *env*, and the length of the whole virus (Fig. 3, Supplementary Table S2). The *vpu* genes of Vpu-type 1 were significantly shorter than those of Vpu-types 2, 3, and 4, differing by an average of 16.4 nucleotide bases. In contrast, the length of the overlap between *env* and *vpu* varied considerably within Vpu-type 1, but remained relatively stable in Vpu-types 2, 3, and 4. Vpu-type 4 showed a consistently stable overlap, with an average length of 82 bases. Vpu-type 3 had the shortest overlap, at 77.5 bases, and Vpu-type 2 had the longest overlap, measuring 89.6 bases. A greater variation in overlap length (± standard deviation) observed in Vpu-types 1 and 3 than in Vpu-types 2 and 4 suggests potential genome instability in the former types. Over time, the overlaps in the Vpu types have stabilized, which is crucial for the virus because overlapping genes is a common strategy by which viruses compact their genomes (Chirico et al., 2010).

**Figure 3.**
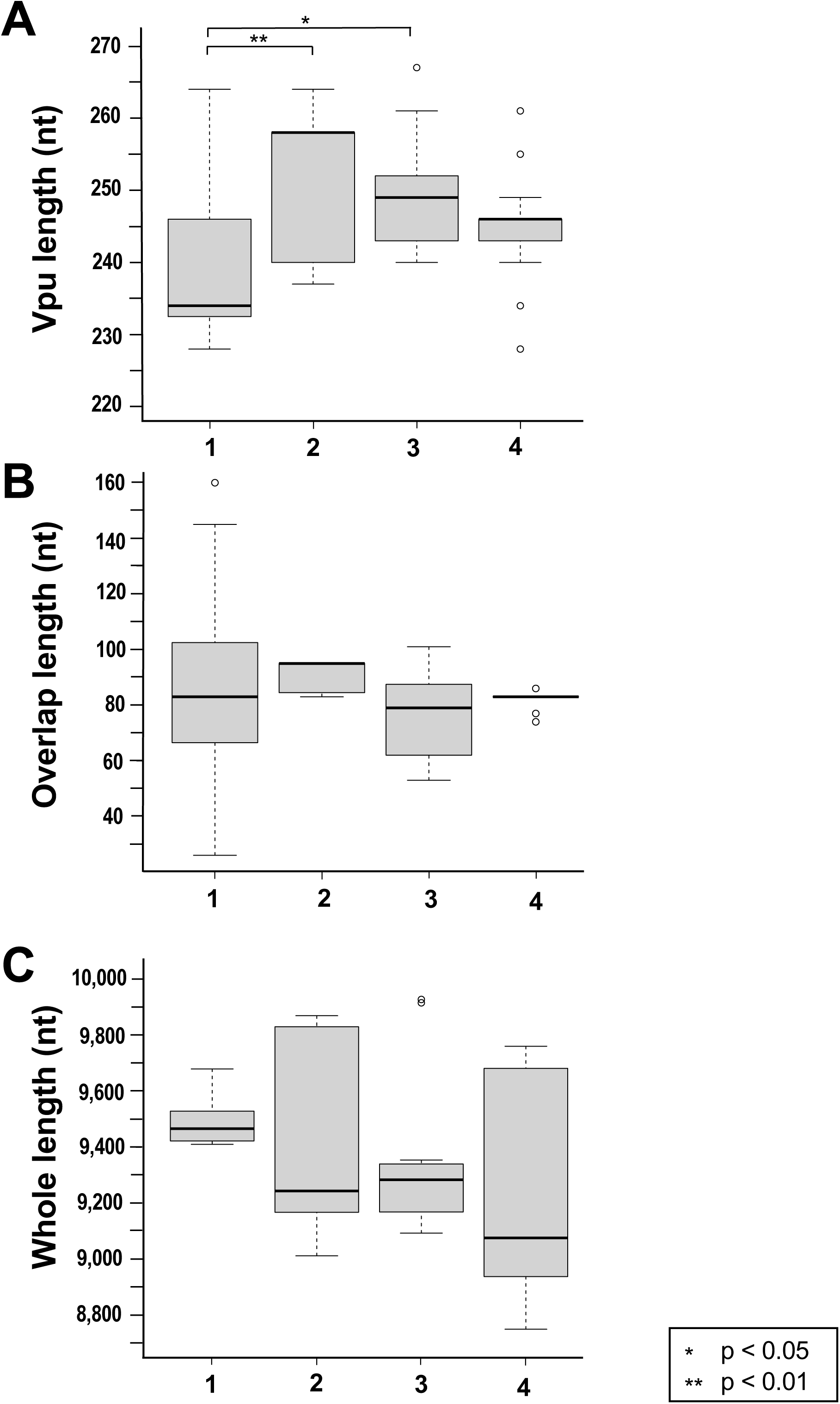
Boxplots comparing the sizes of HIV-1 and SIV *vpu* genes, lengths of overlap, and whole-genome lengths. Boxplots of the nucleotide lengths of *vpu* (A), length of the overlap between *vpu* and *env* (B), and whole-genome lengths (C) are shown (total sequences, n = 56; see Supplementary Table S2). The number below each figure indicates each Vpu-type. Tukey’s method was used to identify significant differences between means (*p < 0.05, **p < 0.01).

Our analysis indicated a trend of decreasing genome sizes across Vpu-types 1‒4. Vpu-type 1 had an average genome length of 9,479.1 bases, Vpu-type 2 had an average length of 9,432.5 bases, Vpu-type 3 had an average length of 9,384.2 bases, and Vpu-type 4 had the shortest length of 9,098.3 bases. Although Vpu-type 1 has a small *vpu* gene, its genome is longer than that of the other types. To gain further insight into the factors contributing to this difference, we compared the lengths of all nine genes in the viral genomes (*gag*, *pol*, *vif*, *tat*, *rev*, *vpu*, *env*, and *nef*). This allowed us to identify specific genomic regions that contribute to the larger genome size of Vpu-type 1 (Supplementary Fig. S3). The comparison of gene lengths suggested that, except for *vpu*, all of the genes in Vpu-type 1 are larger than those in Vpu-type 4. These findings suggest that over the course of evolution, genomic reduction had taken place to create a more efficient virus. It also suggests that the *vpu* gene in Vpu-type 1, the first to acquire the gene, has not undergone significant expansion, explaining its shorter length relative to those of the other Vpu types. Therefore, Vpu-type 1 may be considered to contain the primitive form of *vpu*. Furthermore, whereas Vpu-type 1 primarily uses Vpu to antagonize and downregulate CD4^+^ T cells, SIVcpz uses Nef to antagonize tetherin because SIVcpz Vpu is a very poor tetherin antagonist. Therefore, SIVcpz does not rely on Vpu for both functions. This difference suggests a potential reliance on host-specific adaptation for tetherin antagonism, highlighting the importance of host-specific factors in viral evolution and pathogenesis (Sauter et al., 2009). In contrast, host-specific adaptation may be less critical for the downregulation of CD4^+^ T cells, given the widespread activity of the primate lentiviral Vpu and Nef proteins against human CD4^+^ T cells (Schindler et al., 2006). The transition from utilizing Vpu to Nef to conduct these functions is suggested to have occurred because the use of the *vpu* gene in Vpu-type 1 might not have been favorable for the virus, particularly because these strains were the first to acquire the gene. After the transmission from SIVcpz to HIV-1 group M, HIV-1 group M reverted back to using the Vpu protein to antagonize tetherin, possibly undergoing significant evolutionary adaptations to enhance its antagonistic activities. Among the nonpandemic strains of HIV-1, group N demonstrates a substantial level of anti-tetherin activity, whereas groups O and P do not have this trait. Moreover, group N lacks the ability to downregulate CD4^+^ T cells. These findings allow us to infer that the Vpu protein plays a pivotal role in the pandemic attributed to HIV-1 group M.

### 3.2 Acquisition of Vpu proteins

To clarify how *vpu* was acquired, we constructed a phylogenetic tree of HIV-1, HIV-2, and SIV based on the nucleotide sequence between the *pol* and *env* genes, as indicated in Supplementary Fig. S1. This region was selected due to its high genomic variability and because it contains the *vpu* and *vpx* genes. We retrieved all available, partial and complete HIV and SIV genome sequences from NCBI GenBank. Our initial dataset comprised 10,891 HIV-1, 9,958 HIV-2, and 654 SIV sequences. To maintain the overall diversity of the dataset, we reduced it to 427 HIV-1, 284 HIV-2, and 111 SIV sequences using CD-HIT with a 90% sequence similarity threshold. The midpoint-rooted phylogenetic tree indicated that Vpu-type 1 consists of SIVden, SIVgsn, SIVmon, and SIVmus, which are closely related to the SIVs that infect monkeys, such as the red-tailed monkey (SIVasc), African Sykes’ monkey (SIVsyk), and de Brazza’s monkey (SIVdeb), which lack both the *vpu* and *vpx* genes (Fig. 4). These SIV strains were classified as *vpu*^−^*vpx*^−^ in our study. Specifically, we found a mixture of Vpu-type 1 and *vpu*^−^*vpx*^−^- type sequences of SIVden and SIVdeb, respectively, which suggests that *vpu* arose from the *vpu*^−^*vpx*^−^ type. Because previous studies have reported that the *vpu* gene originated from a common ancestor of SIVgsn and SIVmus (Bibollet-Ruche et al., 2004), we constructed an extensive phylogenetic tree including both HIV and SIV sequences in a novel and comprehensive analysis that delved deeper into the evolutionary history of the *vpu* gene. To validate our findings, an additional tree was constructed using the Pol protein sequences because the *pol* gene is known for its high degree of conservation among the lentiviruses (Jern et al., 2005; Malossi et al., 2020). As on our previous tree, SIVden, SIVmus, and SIVmon were located close to the SIVsyk and SIVasc sequences on the Pol tree, confirming their close relatedness. Moreover, viruses other than SIVsyk, SIVasc, and SIVdeb were positioned more distantly from the other *vpu*^−^*vpx*^−^ viruses, prominently situated near Vpu-type 1. This further supports our findings that Vpu-type 1 is closely related to *vpu*^−^*vpx*^−^ and that Vpu-type 1 contains the oldest and most-primitive form of the *vpu* gene (Supplementary Fig. S4).

**Figure 4.**
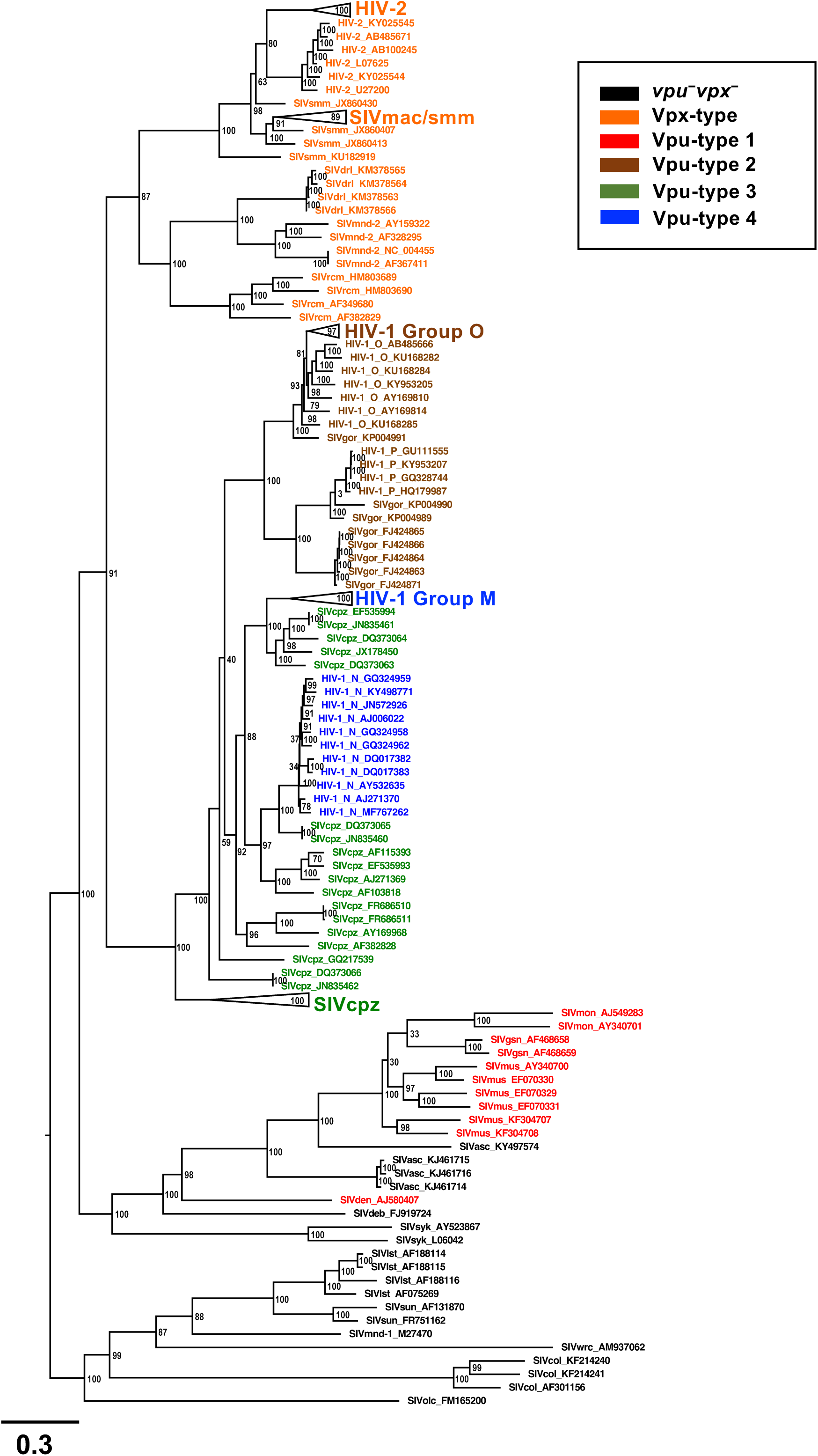
Rooted phylogenetic tree of nucleotide sequences in the *pol–env* region. A midpoint-rooted phylogenetic tree of *pol–env* nucleotide sequences (n = 401; see Supplementary Table S1) was constructed (1,000 bootstrap replicates). The region of focus is shown in Supplementary Fig. S1. Annotations of the viral strains and GenBank IDs, extracted from GenBank, are colored according to the types: *vpu*^−^*vpx*^−^ (black), Vpx-type (orange), Vpu-type 1 (red), Vpu-type 2 (brown), Vpu-type 3 (green), and Vpu-type 4 (blue). SIVmac/smm, SIVcpz, HIV-1 group M, HIV-1 group O, and HIV-2 were collapsed and are shown as black triangles. The scale bar below the tree indicates 0.3 (30%) amino acid substitutions per site, and bootstrap values (1/100), indicating the estimated posterior probabilities, are given at each node.

Thus far, the exact origins of the *vpu* gene remain unclear. To address this question, we performed multiple-sequence alignment of the complete genomes of SIVs (*vpu*^−^*vpx*^−^) and the *vpu* gene. The aim was to identify sequences similar to *vpu* within the SIV genome, on the assumption that the gene originated within the genome itself. We detected several nucleotide sequence similarities in parts of the *vpr* and *env* genes located near the *vpu* gene. As shown in Supplementary Figs. S5 and S6, we specifically compared SIVden *vpu* with *vpr* and *env* of SIVdeb and SIVsyk based on the closeness of the types Vpu-type 1 and *vpu*^−^*vpx*^−^, as the phylogenetic tree shown in Fig. 4. We mapped the positions that shared sequence similarities between the genes onto the structure, indicating their exact positions within the genome. Specifically, we aligned the nucleotide sequences of *vpr* and the first 220 bases of *env* (Supplementary Figs. S5 and S6B) with *vpu*. Notably, in case 1, SIVden *vpu* and SIVdeb *vpr* shared similarities of 46% (75/163) and 62% (73/118) with the 5′ end of *env*, respectively. Conversely, in case 2, comparing SIVden *vpu* with the SIVsyk genome revealed that it shares 45% (68/152) similarity with *vpr* and 43% (44/103) similarity with *env*. Notably, we discovered that the *vpr* and *env* genes in SIVsyk share similarities with the 3′ end of *vpu*. These findings collectively suggest that *vpu* originated through the misalignment of direct repeats within *vpr* and *env*. A replication misalignment, also known as ‘slippage’, may have occurred between the two genes within the SIV genome, leading to the creation of the *vpu* gene (Lovett, 2004). A slipped alignment of the nascent strain with respect to the template may have expanded the directly repeated sequence of *vpr* and *env*, resulting in the formation of a duplicate region. Therefore, the *vpu* gene may be a hybridized form of these two genes. This idea is also supported by the fact that the *vpu* and *env* genes of HIV partially overlap.

### 3.3 Diversification of *vpu*

After the acquisition of *vpu*, the gene underwent a series of evolutionary events detailed in the previous sections, including modifications of its length and the region overlapping the *env* gene, and genome changes—each of which was crucial in shaping the virulent pandemic strain, HIV-1 group M. We constructed an unrooted phylogenetic tree specifically with the Vpu sequences of Vpu-type 4a (HIV-1 group M) to investigate the evolutionary processes and diversification that led to the emergence of group M. We used the same dataset of sequences to construct this tree as used to construct the phylogenetic tree in Fig. 1, but the total number of sequences was reduced to 242. We also included the newly reported strain, variant subtype B (Wymant et al., 2022), in our analysis. We constructed the unrooted phylogenetic tree to clarify the relationship between the viral subtypes and to identify potential differences of Vpu protein among the subtypes. We observed significant divergence within the pre-existing classification of the subtypes. Currently, Vpu-type 4a includes 10 recognized viral subtypes (subtypes A–D, F–H, J–L) (Bbosa et al., 2019; Hemelaar et al., 2006; Sharp & Hahn, 2011). Therefore, in a more specific classification, shown in Fig. 5, we reclassified the previously known subtypes into more detailed categories, which are referred to as ‘protein subtypes’ throughout this study. This reclassification revealed a total of 17 protein subtypes based on their monophyletic clades. Notably, the viral subtypes diverged within their initial classification, suggesting the need for a more thorough and in-depth classification, to allow the comprehensive study of the virus. The Vpu proteins of the subtype B viruses showed particular sequence diversity, and we identified protein subtypes such as BFK-1, which combines viral subtypes B, F, and K that are reflected in its name. Similarly, BD-1 combines subtypes B and D. Interestingly, subtypes A, B, and C showed significant variation within their initial clades, generating three, four, and five newly classified protein subtypes, respectively, within their respective groups. Subtype C showed high divergence, with five monophyletic clades classified as C-1, C-2, C-3, C-4, and C-5. The number of sub-subtypes (or protein subtypes in this case) classified using the full-length genome (nucleotide sequences) differed from the number when using Vpu-type (protein sequences) alone. By contrast, another study identified seven sub-subtypes within subtype A and three within subtype B (Desire et al., 2018), which is fewer than the number we propose for subtype A and more than the number we propose for subtype B. However, the grouping of subtype B and D together is consistent in both studies, supporting our new classification. This concurrence validates our new classification of these subtypes and underscores the importance of reclassifying the subtypes to better detect and trace the epidemiological changes in the virus. Our analysis also highlights the misannotations in the existing sequence data, revealing instances where certain subtypes have been erroneously intermixed within clusters due to annotation errors.

**Figure 5.**
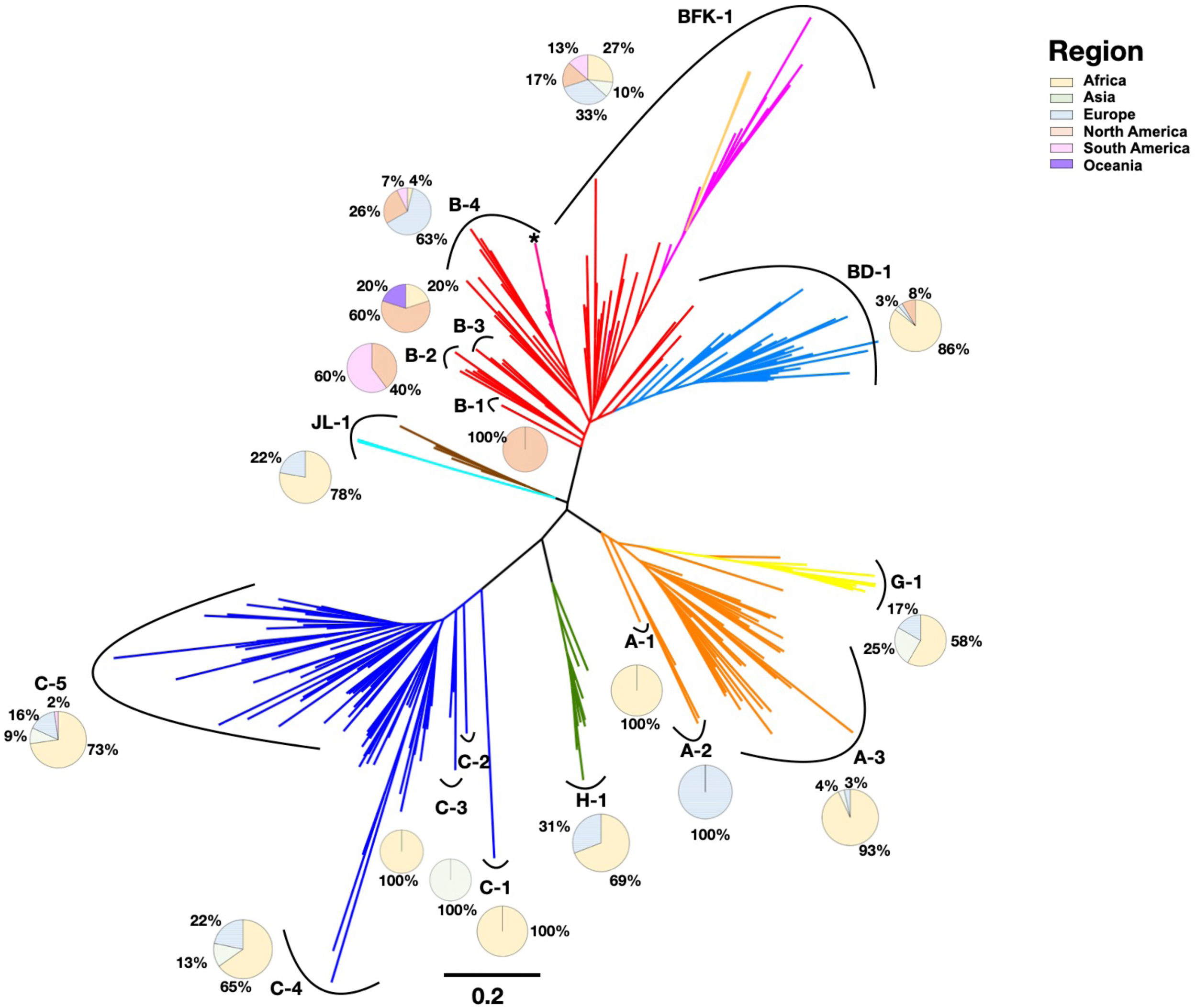
Unrooted phylogenetic tree of Vpu proteins showing the regional distributions of the viruses. The unrooted phylogenetic tree was constructed from Vpu protein sequences (1,000 bootstrap replicates) (n = 242; see Supplementary Table S3). The branches of the phylogenetic tree are colored according to their official viral subtypes. The pie chart shows the regional prevalence of the protein subtypes, using the following colors: Asia (light green), Africa (yellow), Oceania (purple), Europe (light blue), North America (light orange), and South America (pink). The scale bar below the tree indicates 0.2 (20%) amino acid substitutions per site and bootstrap values (1/100), indicating estimated posterior probabilities, are given at each node.

To identify protein-subtype-specific (newly annotated types) differences and subtype-specific differences in these strains, we investigated possible differences in their regional and yearly prevalence (Fig. 5 and Supplementary Fig. S7). We compiled a table of prevalence data for regions and years in decadal intervals (Supplementary Table S3) from the sequence annotations of the subtypes. We then generated a pie chart to visually represent the data. Consequently, we identified temporal (decadal) and regional differences between the protein subtypes. For instance, variations in the years of prevalence were observed within types B-1, B-2, B-3, and B-4. Specifically, B-1 included sequences from 1990– 99, whereas for B-2, 75% of sequences were from 2010–19 and 25% from 2000–09. Moreover, in B-3, 50% of sequences were from 2000–09, 25% from 1980–89, and 25% from 2010–19. Lastly, in B-4, 81% of sequences were from 2000–09, 11% from 1990–99, 4% from 2010–19, and 4% from 1980–89. The distributions of regions of prevalence also showed variations. For instance, B-1 consisted solely of sequences from North America. By contrast, B-2 displayed a mix of sequences from South America (60%) and North America (40%). In B-3, 60% of sequences were from North America, 20% from Oceania, and another 20% from Africa. Lastly, in B-4, 63% of sequences were from Europe, 26% from North America, 7% from South America, and 4% from Africa (Supplementary Table S4). We also incorporated the Vpu proteins of 135 circulating recombinant forms (1–135) together with those of HIV-1 group M (Vpu-type 4a) and constructed another unrooted phylogenetic tree to visualize the diversification of the protein within HIV-1 (Supplementary Fig. S8, Supplementary Table S5). We identified an additional distinct protein group, designated ‘R1’ to represent recombinants that were clustered together. The distinct clustering of R1 indicates that the Vpu protein is evolving separately from other protein subtypes, highlighting the continuous evolution of Vpu. R1 is predominantly composed of sequences collected from Asia in 2010–19. We conducted a multiple sequence analysis between neighboring protein subtypes G-1 and A-4, which revealed that R1 contains sequences derived from these protein subtypes (Supplementary Fig. S9). Therefore, a total of 18 (17 protein subtypes + 1 recombinant group R1) protein subtypes were classified overall just within HIV-1 group M. This finding suggests the potential evolutionary relationships between the identified groups and the high divergence of the recombinants, and highlights the ongoing, rapid evolution of the *vpu* gene in the viral subtypes and the circulating recombinant forms.

A variant subtype B strain that is highly virulent and reduces CD4 cells twice as efficiently as the normal subtype B strain has recently been reported (Wymant et al., 2022). To visualize its evolutionary lineage, we constructed a rooted phylogenetic tree of Vpu types including this variant strain B from Vpu-type 4a, created a multiple-sequence alignment, and used multi-Harmony to identify specific differences between the *vpu* genes of subtype B and the virulent variant B subtype (Supplementary Figs. S10 and S11). The tree revealed that the variant B strain is still evolving because it is located at the very top of the tree, branching from subtype B. The multiple-sequence alignment of the amino acids of Vpu from the two strains showed that the variant is more highly conserved than the common subtype. Notably, positions such as 11 (residue 8), 19 (residue 16), 50 (residue 47), 76 (residue 73), and 80 (residue 77) in the variant B subtype showed more than 94% conservation within the same strain, whereas subtype B showed a significantly lower conservation rates within the same strain. The comparison of the two strains at these five positions showed high variation rates of 81%, 65%, 59%, 82%, and 71%, respectively. These observations suggest that even minor amino acid changes within Vpu play crucial roles in influencing the virulence of HIV-1.

## 4. Conclusions

*vpu* is a key gene in the HIV-1 genome and in the genomes of SIVs, such as SIVcpz and SIVmus. By performing a comprehensive analysis, including bioinformatic approaches comprising sequence analyses and phylogenetics, we traced the evolutionary history of *vpu* and developed a tentative evolutionary model of the gene, outlining the steps of its acquisition and evolution (Fig. 6). It is important to note that this study primarily presents a working hypothesis because there are inherent limitations to relying solely on computational methods. First, we propose that the acquisition of the gene occurred in monkeys, such as the mona monkey (SIVmon), moustached guenon (SIVmus), and greater spot-nosed monkey (SIVgsn) from African Sykes’ monkey (SIVsyk) and red-tailed monkey (SIVasc), which are *vpu*^−^*vpx*^−^ SIVs, through a misalignment duplication. Second, the gene underwent phases of trial-and-error changes in the genome, the length of *vpu*, and the length of the overlap between *vpu* and *env*, possibly to allow the virus to replicate more efficiently, and consequently creating a more infectious virus. In the latter part of our study, we focused exclusively on the evolution of *vpu* in HIV-1 group M (Vpu-type 4a). We demonstrated that *vpu* diversified rapidly over time and reclassified the viral subtypes and circulating recombinants into 18 (17 protein subtypes + 1 recombinant type) more-specific groups. This highlights the importance of the detailed classifications of both viral subtypes and circulating recombinant forms in extending our understanding of the evolutionary patterns of the virus. We also examined regional and temporal (decadal) trends in the protein subtypes and identified variations in prevalence between and within the protein types.

**Figure 6.**
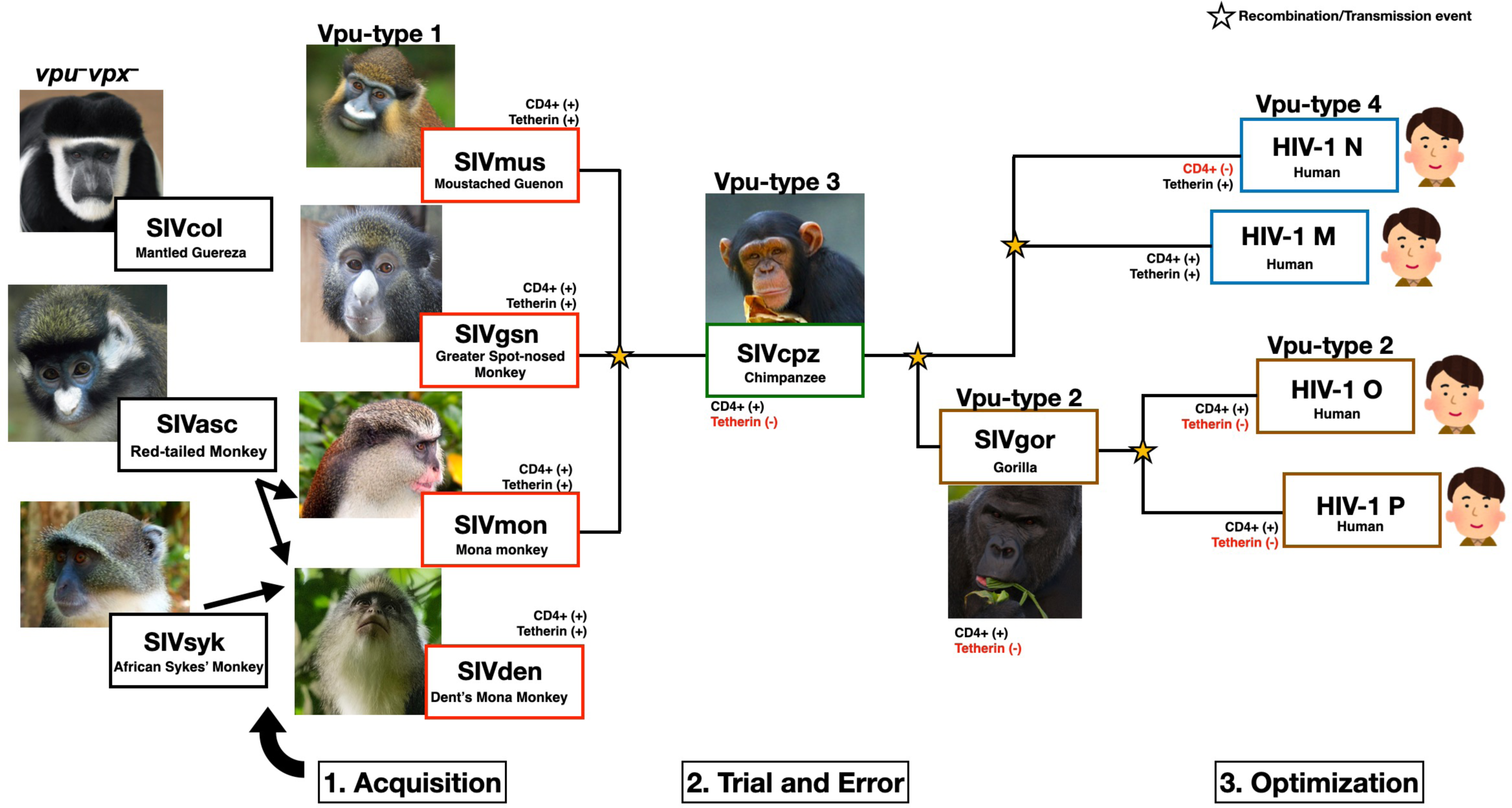
Proposed model of *vpu* gene evolution in SIV and HIV-1 strains. Schematic representation of the model of *vpu* evolution. The types are labeled above each picture. The viral strain and name of each monkey species are written in the individually colored boxes. Black arrows indicate the possible ancestors of SIVmon and SIVden. Vpu evolved through phases of trial-and-error optimization to create an ideal gene. Yellow stars indicate recombination and transmission events. Pictures were taken from creative commons (see Materials and methods).

Lastly, we compared the *vpu* sequences of variant subtype B, a particularly virulent strain, with that of the normal common subtype B. The *vpu* gene sequence is more highly conserved in the variant strain than in the common strain. The variations observed in the transmembrane and cytoplasmic domains may influence the functions of the encoded protein. Although the changes are subtle, they could play a pivotal role in the protein’s function, potentially contributing to the increased virulence of the variant strain. Although we speculated a potential correlation between the Vpu protein and the decline in CD4 cell levels, it is crucial to acknowledge that no experimental studies of the *vpu* gene of the new variant have yet been conducted. Nevertheless, our methodological approach may be useful in determining the evolution of the virus. By understanding how viruses evolve, we can track the emergence of new strains and monitor their spread. This will provide valuable insights into their adaptation and evolution over time that is particularly critical for pathogenic viruses with high mutation rates, such as HIV and severe acute respiratory syndrome coronavirus 2 (SARS-CoV-2). Such understanding is also crucial for developing effective strategies to combat viral diseases and mitigate their impact on public health.

## Data availability

The data supporting the findings of this work are available within the paper and its Supplementary Information files.

## Acknowledgments

We thank all the members of the RNA Group at the Institute for Advanced Biosciences of Keio University, Japan, for their insightful discussions. We also thank Miura Masahiro and Phillip Yamamoto for helping to create the Python script required for this study. This work was supported, in part, by research funds from the Yamagata Prefectural Government and Tsuruoka City, Japan. The funding bodies played no role in the study design, data collection or analysis, the decision to publish, or the preparation of the manuscript.

