## Supplementary file for "Possible acquisition and molecular evolution of *vpu* genes inferred from comprehensive sequence analysis of human and simian immunodeficiency viruses"

**Supplementary Table S1.**

| | | No. nucleotide sequences (pol-env) | | No. Vpu protein sequences | | | No. Pol protein sequences | | |
|---|---|---|---|---|---|---|---|---|---|
| **Types** | **Host** | **No. of sequences** | **After CD-HIT** | **No. of sequence** | **After CD-HIT** | **Removal Of Xs** | **No. of sequence** | **After CD-HIT** | **Removal Of Xs** |
| **Prototype** | **SIVagm** | 0 | 0 | - | - | - | 3 | 3 | 3 |
| | **SIVasc** | 3 | 1 | - | - | - | 1 | 1 | 1 |
| | **SIVcol** | 3 | 3 | - | - | - | 0 | 0 | 0 |
| | **SIVsun** | 2 | 2 | - | - | - | 1 | 1 | 1 |
| | **SIVsyk** | 3 | 3 | - | - | - | 1 | 1 | 1 |
| | **SIVmnd-1** | 4 | 3 | - | - | - | 0 | 0 | 0 |
| | **SIVolc** | 1 | 1 | - | - | - | 0 | 0 | 0 |
| | **SIVlst** | 1 | 1 | - | - | - | 1 | 1 | 0 |
| | **SIVwrc** | 3 | 3 | - | - | - | 2 | 2 | 2 |
| **Vpx-type** | **HIV-2** | 55 | 12 | - | - | - | 42 | 42 | 21 |
| | **SIVdrl** | 5 | 1 | - | - | - | 0 | 0 | 0 |
| | **SIVmac** | 527 | 5 | - | - | - | 1 | 1 | 1 |
| | **SIVmnd-2** | 4 | 3 | - | - | - | 2 | 2 | 1 |
| | **SIVsmm** | 47 | 9 | - | - | - | 7 | 7 | 3 |
| | **SIVrcm** | 4 | 4 | - | - | - | 0 | 0 | 0 |
| **Vpu-type** | **IV-1 Group** | 9,931 | 296 | 38,708 | 131 | 105 | 19,873 | 94 | 64 |
| | **Group N** | 11 | 2 | 12 | 5 | 5 | 6 | 1 | 0 |
| | **Group O** | 55 | 16 | 69 | 14 | 14 | 3 | 1 | 1 |
| | **Group P** | 4 | 1 | 3 | 1 | 1 | 2 | 0 | 0 |
| | **SIVcpz** | 209 | 20 | 36 | 5 | 5 | 9 | 7 | 5 |
| | **SIVgor** | 8 | 4 | 4 | 3 | 2 | 3 | 1 | 0 |
| | **SIVden** | 1 | 1 | 1 | 1 | 1 | 1 | 1 | 1 |
| | **SIVgsn** | 2 | 2 | 1 | 1 | 1 | 2 | 1 | 1 |
| | **SIVmon** | 2 | 2 | 1 | 2 | 1 | 1 | 1 | 1 |
| | **SIVmus** | 6 | 6 | 6 | 6 | 6 | 4 | 4 | 4 |
| **Total** | | 10,891 | 401 | 38,841 | 169 | 141 | 19,965 | 172 | 111 |

**Supplementary Table S1.** Summary of the viral sequences used to construct the phylogenetic trees.

CD-HIT: a program used for clustering and comparing protein or nucleotide sequences (Fu et al., 2012). ‘X’ represents an unknown amino acid; these were removed in our study Xs’:an unknown amino acid.

**Supplementary Table S2:**

| Vpu-type | Vpu length (nt) | Overlap length (nt) | Genome length (nt) |
|---|---|---|---|
| **Type 1 (n=11)** | 239.5 ± 10.7 | 88.5 ± 38.5 | 9490.3 ± 90.3* |
| **Type 2 (n=12)** | 252.0 ± 10.5 | 90.5 ± 5.6 | 9432.5 ± 351.6 |
| **Type 3 (n=12)** | 249.5 ± 8.2 | 77.1 ± 14.8 | 9348.4 ± 279.9 |
| **Type 4 (n=21)** | 246.0 ± 7.3 | 81.7 ± 3.2 | 9207.7 ± 355.4 |

**Supplementary Table S2.** Summary of the nucleotide sequence lengths of vpu, the lengths of vpu–env overlapping region, and whole-genome lengths of the virus. For each sample size, the mean and standard deviation of the lengths per category are provided. Due to the limited availability of Vpu-type 1 sequences, we used all the available sequences, including partial genome sequences.

**Supplementary Table S3:**

**A**

| **Subtype** | **Africa** | **Asia** | **Europe** | **North America** | **South America** | **Oceania** | **NA** |
|---|---|---|---|---|---|---|---|
| **A** | 28 | 2 | 4 | 0 | 0 | 0 | 0 |
| **B** | 0 | 4 | 5 | 21 | 7 | 1 | 1 |
| **C** | 51 | 8 | 12 | 0 | 1 | 0 | 0 |
| **D** | 33 | 0 | 0 | 0 | 0 | 0 | 1 |
| **F** | 4 | 0 | 6 | 0 | 2 | 0 | 0 |
| **G** | 6 | 2 | 2 | 0 | 0 | 0 | 0 |
| **H** | 9 | 0 | 4 | 0 | 0 | 0 | 0 |
| **J** | 4 | 0 | 2 | 0 | 0 | 0 | 0 |
| **K** | 2 | 0 | 0 | 0 | 0 | 0 | 0 |
| **L** | 3 | 0 | 0 | 0 | 0 | 0 | 0 |
| **VB** | 0 | 0 | 17 | 0 | 0 | 0 | 0 |
| **Total** | 140 | 16 | 52 | 21 | 10 | 1 | 2 |

**B**

| **Subtype** | **1980-1989** | **1990-1999** | **2000-2009** | **2010-2019** | **2020-2022** | **NA** |
|---|---|---|---|---|---|---|
| **A** | 1 | 4 | 18 | 10 | 0 | 0 |
| **B** | 2 | 4 | 14 | 13 | 0 | 7 |
| **C** | 0 | 2 | 36 | 31 | 0 | 3 |
| **D** | 1 | 10 | 13 | 10 | 1 | 0 |
| **F** | 0 | 2 | 4 | 6 | 0 | 0 |
| **G** | 0 | 2 | 2 | 5 | 0 | 1 |
| **H** | 0 | 6 | 7 | 0 | 0 | 0 |
| **J** | 0 | 4 | 2 | 0 | 0 | 0 |
| **K** | 0 | 2 | 0 | 0 | 0 | 0 |
| **L** | 1 | 1 | 0 | 0 | 0 | 0 |
| **VB** | 0 | 0 | 17 | 0 | 0 | 0 |
| **Total** | 5 | 37 | 113 | 75 | 1 | 11 |

**Supplementary Table S3.** Summary of the sequences used to construct the phylogenetic tree of HIV-1 group M Vpu, categorized by geographic and temporal (decadal) breakdowns. The number of sequences is categorized according to the subtype. VB, new variant B subtype.

**Supplementary Table S4:**

**A**

| Protein sub-type | Percentage (%) | | | | | |
|---|---|---|---|---|---|---|
| | Africa | Asia | Europe | North America | South America | Oceania |
| A-1 | 100 | - | - | - | - | - |
| A-2 | - | - | 100 | - | - | - |
| A-3 | 93 | 4 | 3 | - | - | - |
| AG-1 | 58 | 25 | 17 | - | - | - |
| B-1 | - | - | - | 100 | - | - |
| B-2 | - | - | - | 40 | 60 | - |
| B-3 | 20 | - | - | 60 | - | 20 |
| B-4 | 4 | - | 63 | 26 | 7 | - |
| BD-1 | 86 | 3 | 3 | 8 | - | - |
| BFK | 27 | 10 | 33 | 17 | 13 | - |
| C-1 | 100 | - | - | - | - | - |
| C-2 | - | 100 | - | - | - | - |
| C-3 | 100 | - | - | - | - | - |
| C-4 | 65 | 13 | 22 | - | - | - |
| C-5 | 73 | 9 | 16 | - | 2 | - |
| H-1 | 69 | - | 31 | - | - | - |
| JL-1 | 78 | - | 22 | - | - | - |

**B**

| Protein sub-type | Percentage (%) | | | | |
|---|---|---|---|---|---|
| | 1980-1989 | 1990-1999 | 2000-2009 | 2010-2019 | 2020-2022 |
| A-1 | - | - | 100 | - | - |
| A-2 | - | - | 33 | 67 | - |
| A-3 | 3 | 14 | 55 | 28 | - |
| AG-1 | - | 30 | 30 | 50 | - |
| B-1 | - | 100 | - | - | - |
| B-2 | - | - | 25 | 75 | - |
| B-3 | 25 | - | 50 | 25 | - |
| B-4 | 4 | 11 | 81 | 4 | - |
| BD-1 | 3 | 24 | 33 | 37 | 3 |
| BFK | - | 15 | 39 | 46 | - |
| C-1 | - | - | 100 | - | - |
| C-2 | - | - | - | 100 | - |
| C-3 | - | - | 67 | 33 | - |
| C-4 | - | - | 36 | 64 | - |
| C-5 | - | 5 | 59 | 36 | - |
| H-1 | - | 46 | 54 | - | - |
| JL-1 | 11 | 56 | 33 | - | - |

**Supplementary Table S4.** Summary of the regional and temporal (decadal) prevalence of the newly classified protein subtypes. (A) Geographical and (B) temporal (decadal) breakdown prevalence of the respective protein subtypes (percentages). The prevalence of each protein subtype is represented in the accompanying pie chart (see Fig. 5 and Supplementary Fig. S7) The temporal data are organized into decades.

**Supplementary Table S5:**

**A**

| CRF | Africa | Asia | Europe | North America | South America | Oceania | NA |
|---|---|---|---|---|---|---|---|
| 15 | 0 | 1 | 0 | 0 | 0 | 0 | 0 |
| 22 | 1 | 0 | 0 | 0 | 0 | 0 | 0 |
| 33 | 0 | 1 | 0 | 0 | 0 | 0 | 0 |
| 34 | 0 | 1 | 0 | 0 | 0 | 0 | 0 |
| 48 | 0 | 1 | 0 | 0 | 0 | 0 | 0 |
| 52 | 0 | 1 | 0 | 0 | 0 | 0 | 0 |
| 53 | 0 | 1 | 0 | 0 | 0 | 0 | 0 |
| 54 | 0 | 1 | 0 | 0 | 0 | 0 | 0 |
| 55 | 0 | 1 | 0 | 0 | 0 | 0 | 0 |
| 58 | 0 | 1 | 0 | 0 | 0 | 0 | 0 |
| 67 | 0 | 1 | 0 | 0 | 0 | 0 | 0 |
| 68 | 0 | 1 | 0 | 0 | 0 | 0 | 0 |
| 69 | 0 | 1 | 0 | 0 | 0 | 0 | 0 |
| 76 | 0 | 1 | 0 | 0 | 0 | 0 | 0 |
| 77 | 0 | 1 | 0 | 0 | 0 | 0 | 0 |
| 78 | 0 | 1 | 0 | 0 | 0 | 0 | 0 |
| 79 | 0 | 1 | 0 | 0 | 0 | 0 | 0 |
| 80 | 0 | 1 | 0 | 0 | 0 | 0 | 0 |
| 82 | 0 | 1 | 0 | 0 | 0 | 0 | 0 |
| 83 | 0 | 1 | 0 | 0 | 0 | 0 | 0 |
| 96 | 0 | 1 | 0 | 0 | 0 | 0 | 0 |
| 97 | 0 | 1 | 0 | 0 | 0 | 0 | 0 |
| 100 | 0 | 1 | 0 | 0 | 0 | 0 | 0 |
| 101 | 0 | 1 | 0 | 0 | 0 | 0 | 0 |
| 102 | 0 | 1 | 0 | 0 | 0 | 0 | 0 |
| 103 | 0 | 1 | 0 | 0 | 0 | 0 | 0 |
| 104 | 0 | 1 | 0 | 0 | 0 | 0 | 0 |
| 105 | 0 | 1 | 0 | 0 | 0 | 0 | 0 |
| 106 | 0 | 1 | 0 | 0 | 0 | 0 | 0 |
| 109 | 0 | 1 | 0 | 0 | 0 | 0 | 0 |
| 111 | 0 | 1 | 0 | 0 | 0 | 0 | 0 |
| 112 | 0 | 1 | 0 | 0 | 0 | 0 | 0 |
| 113 | 0 | 1 | 0 | 0 | 0 | 0 | 0 |
| 114 | 0 | 1 | 0 | 0 | 0 | 0 | 0 |
| 115 | 0 | 1 | 0 | 0 | 0 | 0 | 0 |
| 116 | 0 | 1 | 0 | 0 | 0 | 0 | 0 |
| 117 | 0 | 1 | 0 | 0 | 0 | 0 | 0 |
| 121 | 0 | 1 | 0 | 0 | 0 | 0 | 0 |
| 125 | 0 | 1 | 0 | 0 | 0 | 0 | 0 |
| 126 | 0 | 1 | 0 | 0 | 0 | 0 | 0 |
| 137 | 0 | 1 | 0 | 0 | 0 | 0 | 0 |
| 138 | 0 | 1 | 0 | 0 | 0 | 0 | 0 |
| 140 | 0 | 1 | 0 | 0 | 0 | 0 | 0 |
| 143 | 0 | 1 | 0 | 0 | 0 | 0 | 0 |
| **Total** | 1 | 43 | 0 | 0 | 0 | 0 | 0 |

**B**

| CRF | 1980-1989 | 1990-1999 | 2000-2009 | 2010-2019 | 2020-2022 | NA |
|---|---|---|---|---|---|---|
| 15 | 0 | 0 | 1 | 0 | 0 | 0 |
| 22 | 0 | 0 | 1 | 0 | 0 | 0 |
| 33 | 0 | 0 | 1 | 0 | 0 | 0 |
| 34 | 0 | 1 | 0 | 0 | 0 | 0 |
| 48 | 0 | 0 | 1 | 0 | 0 | 0 |
| 52 | 0 | 1 | 0 | 0 | 0 | 0 |
| 53 | 0 | 0 | 0 | 1 | 0 | 0 |
| 54 | 0 | 0 | 1 | 0 | 0 | 0 |
| 55 | 0 | 0 | 0 | 1 | 0 | 0 |
| 58 | 0 | 0 | 0 | 1 | 0 | 0 |
| 67 | 0 | 0 | 0 | 1 | 0 | 0 |
| 68 | 0 | 0 | 0 | 1 | 0 | 0 |
| 69 | 0 | 0 | 1 | 0 | 0 | 0 |
| 76 | 0 | 0 | 0 | 1 | 0 | 0 |
| 77 | 0 | 0 | 0 | 1 | 0 | 0 |
| 78 | 0 | 0 | 0 | 1 | 0 | 0 |
| 79 | 0 | 0 | 0 | 1 | 0 | 0 |
| 80 | 0 | 0 | 0 | 1 | 0 | 0 |
| 82 | 0 | 0 | 0 | 1 | 0 | 0 |
| 83 | 0 | 0 | 0 | 1 | 0 | 0 |
| 96 | 0 | 0 | 0 | 1 | 0 | 0 |
| 97 | 0 | 0 | 0 | 1 | 0 | 0 |
| 100 | 0 | 0 | 0 | 1 | 0 | 0 |
| 101 | 0 | 0 | 0 | 1 | 0 | 0 |
| 102 | 0 | 0 | 0 | 1 | 0 | 0 |
| 103 | 0 | 0 | 0 | 1 | 0 | 0 |
| 104 | 0 | 0 | 0 | 1 | 0 | 0 |
| 105 | 0 | 0 | 0 | 1 | 0 | 0 |
| 106 | 0 | 0 | 0 | 1 | 0 | 0 |
| 109 | 0 | 0 | 0 | 1 | 0 | 0 |
| 111 | 0 | 0 | 0 | 1 | 0 | 0 |
| 112 | 0 | 0 | 0 | 1 | 0 | 0 |
| 113 | 0 | 0 | 0 | 1 | 0 | 0 |
| 114 | 0 | 0 | 0 | 1 | 0 | 0 |
| 115 | 0 | 0 | 0 | 1 | 0 | 0 |
| 116 | 0 | 0 | 0 | 1 | 0 | 0 |
| 117 | 0 | 0 | 0 | 1 | 0 | 0 |
| 121 | 0 | 0 | 0 | 1 | 0 | 0 |
| 125 | 0 | 0 | 0 | 1 | 0 | 0 |
| 126 | 0 | 0 | 0 | 0 | 1 | 0 |
| 137 | 0 | 0 | 0 | 1 | 0 | 0 |
| 138 | 0 | 0 | 0 | 1 | 0 | 0 |
| 140 | 0 | 0 | 0 | 0 | 1 | 0 |
| 143 | 0 | 0 | 0 | 0 | 1 | 0 |
| **Total** | 0 | 2 | 6 | 33 | 3 | 0 |

**Supplementary Table S5.** Summary of the regional and temporal (decadal) prevalence of R1. (A) Geographic and (B) temporal breakdown of the prevalence of circulating recombinant forms (CRFs) in the R1 cluster. The temporal data are organized into decades.

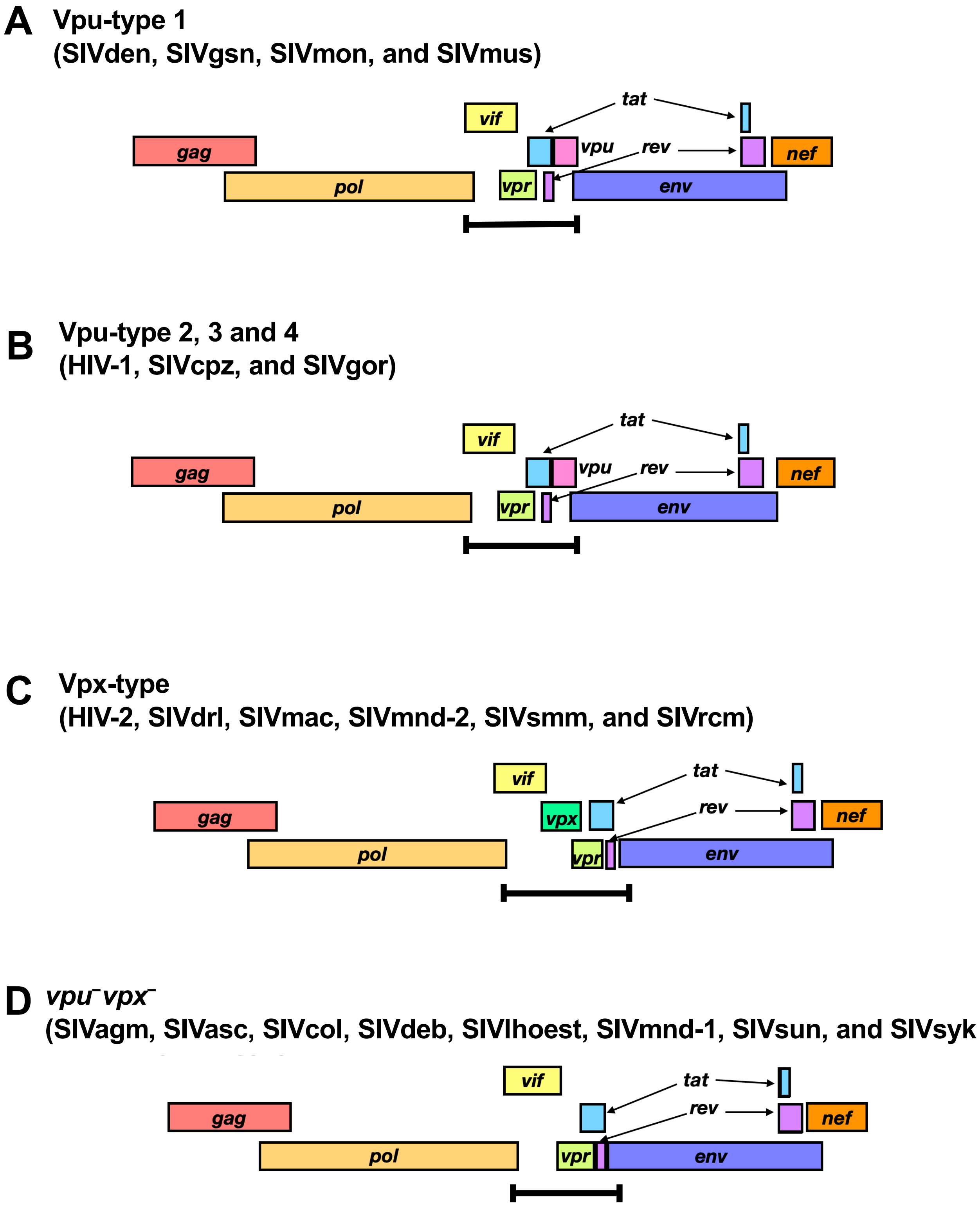


**Supplementary Figure S1.** Schematic representation of the viral genome structures based on the types of viral accessory genes. The genome organization of the Vpu-types are shown, with color-coded genes. Green and light pink indicates *vpx* and *vpu*, respectively, and are only found in certain types. (A) Vpu-type 1 includes SIVden, SIVgsn, SIVmon, and SIVmus. (B) Vpu-types 2, 3, and 4 include HIV-1 (groups M, N, O and P), SIVcpz, and SIVgor. (C) Vpx-type includes HIV-2, SIVdrl, SIVmac, SIVmnd-2, SIVsmm, and SIVrcm. (D) *vpu*⁻*vpx*⁻ includes SIVagm, SIVasc, SIVcol, SIVdeb, SIVlhoest, SIVmnd-1, SIVsun, and SIVsyk.

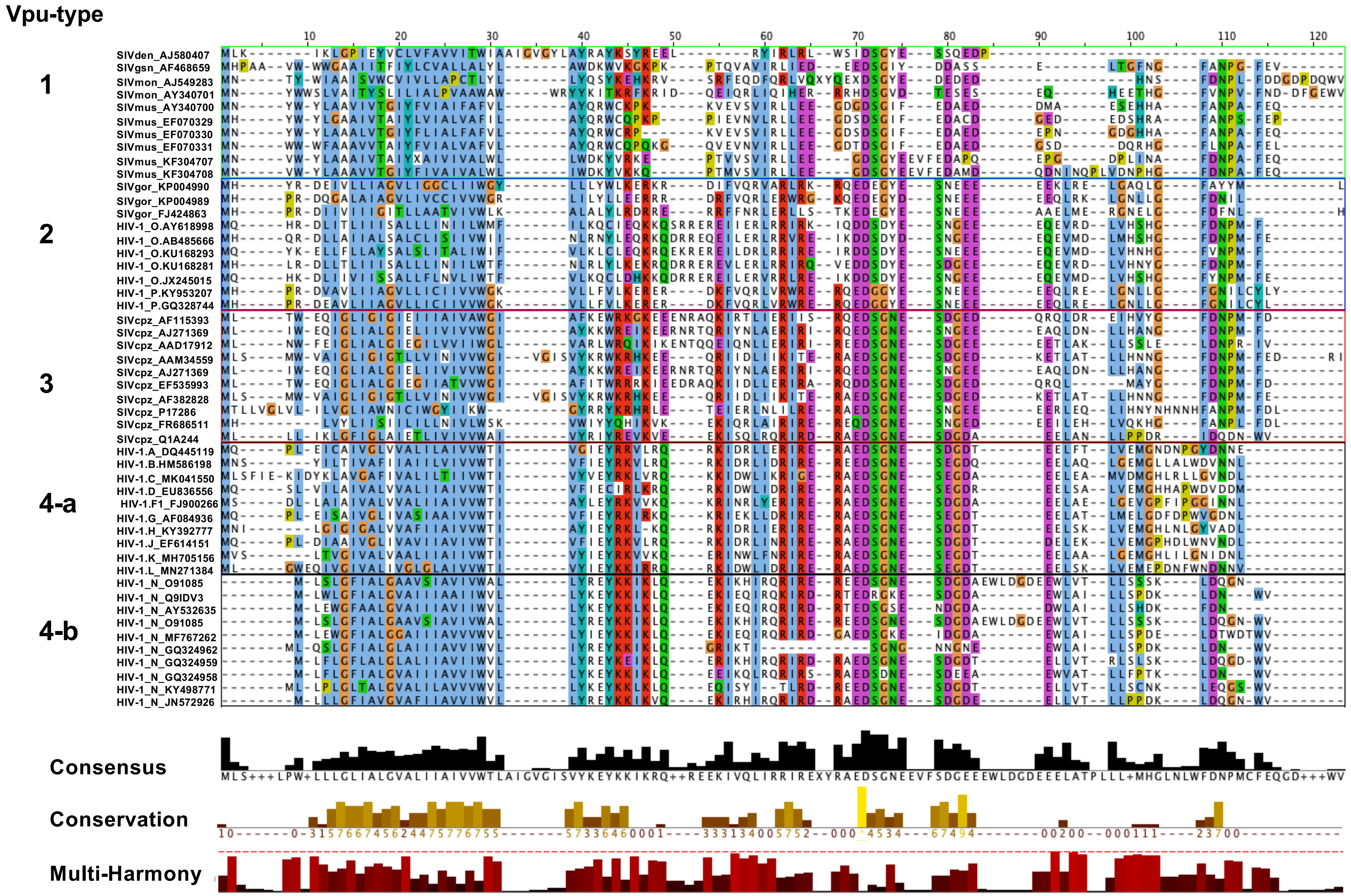


**Supplementary Figure S2.** Multiple-amino-acid-sequence alignment of Vpu proteins. Multiple-sequence alignment of each type of Vpu protein, colored according to the ClustalX color scheme (n = 10). The name of the virus with the corresponding Vpu protein and GenBank ID of the protein are shown along the alignments. The conservation histogram below the alignment describes the sequence conservation of the residues according to their physiochemical properties. The quality annotation represents the likelihood of observing a mutation in any particular column of the alignment based on the BLOSUM62 matrix scores. The consensus histogram shows the conserved regions. The nonconserved residues are marked with '+'. Multi-Harmony histogram shows subtype-specific residues. Transmembrane domain and cytoplasmic domain in Vpu protein are shown above the alignment, with their positions.

**A**

| Vpu-type | *gag* | *pol* | *vif* |
|---|---|---|---|
| **Type 1 (n=8)** | 1554.8 ± 50.9 | 3140.0 ± 286.3 | 731.3 ± 28.0 |
| **Type 2 (n=12)** | 1490.3 ± 11.8 | 3039.8 ± 27.9 | 581.3 ± 5.6 |
| **Type 3 (n=12)** | 1530.8 ± 35.2 | 3016.5 ± 17.8 | 585.5 ± 8.0 |
| **Type 4 (n=12)** | 1500.0 ± 17.8 | 3015.1 ± 15.1 | 579.4 ± 2.0 |
| **Vpu-type** | ***vpr*** | ***tat*** | ***rev*** |
| **Type 1 (n=8)** | 399.0 ± 23.1 | 363.0 ± 60.2 | 357.8 ± 50.5 |
| **Type 2 (n=12)** | 301.0 ± 3.0 | 276.3 ± 54.4 | 313.0 ± 2.0 |
| **Type 3 (n=12)** | 280.5 ± 17.3 | 333.5 ± 50.1 | 345.0 ± 30.5 |
| **Type 4 (n=12)** | 294.5 ± 16.4 | 301.7 ± 18.8 | 348.0 ± 19.5 |
| **Vpu-type** | ***vpu*** | ***env*** | ***nef*** |
| **Type 1 (n=8)** | 238.1 ± 11.8 | 2648.3 ± 35.8 | 670.1 ± 32.0 |
| **Type 2 (n=12)** | 252.0 ± 10.5 | 2631.5 ± 37.8 | 635.3 ± 4.3 |
| **Type 3 (n=12)** | 249.5 ± 8.2 | 2587.3 ± 65.1 | 610.8 ± 15.3 |
| **Type 4 (n=12)** | 246.0 ± 7.3 | 2574.9 ± 56.8 | 628.2 ± 10.7 |

**B**

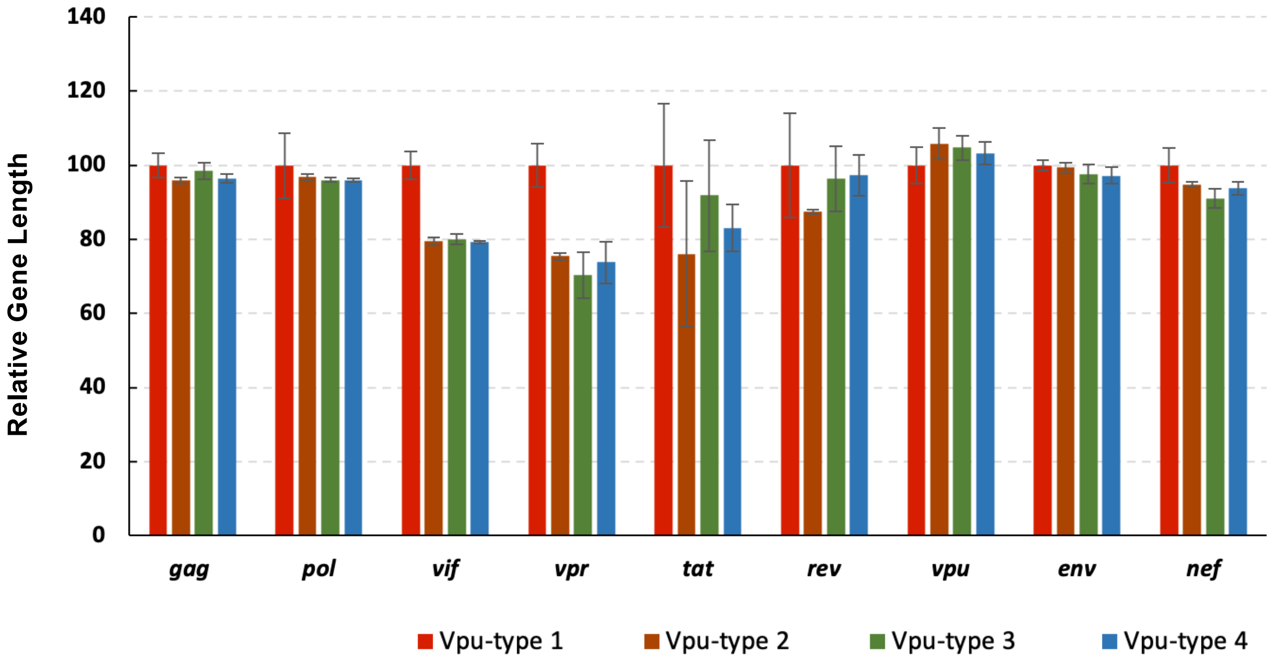


**Supplementary Figure S3.** Size comparisons of the lentiviral genes in the Vpu-types. (A) Average nucleotide sequence lengths of genes *gag*, *pol*, *vif*, *vpr*, *tat*, *rev*, *vpu*, *env*, and *nef*. (B) Normalized bar plot showing gene lengths relative to Vpu-type 1. Bar plots are colored according to type: Vpu-type 1 (red), Vpu-type 2 (brown), Vpu-type 3 (green), and Vpu-type 4 (red).

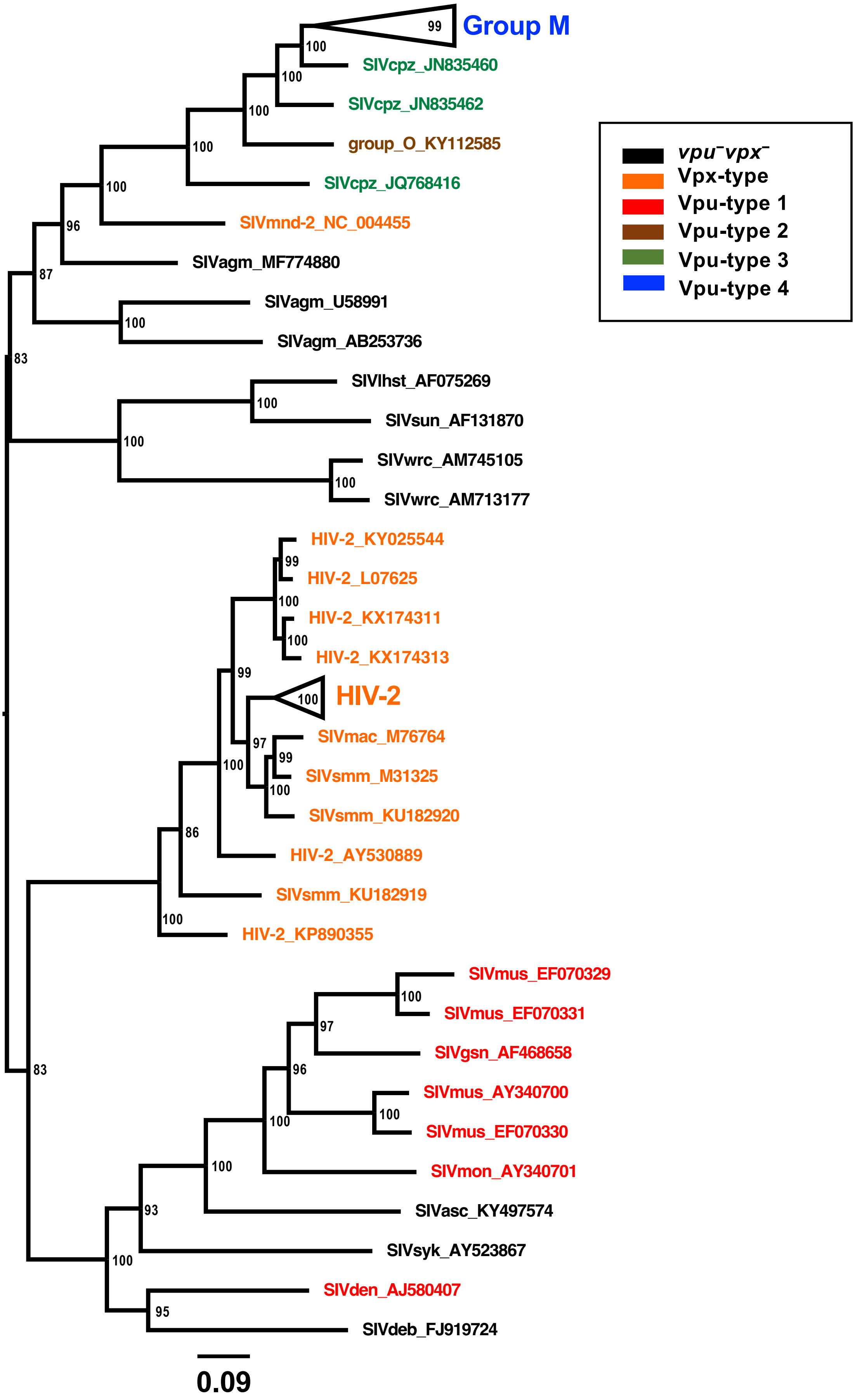


**Supplementary Figure S4.** Rooted phylogenetic tree of Pol protein sequences. A midpoint-rooted phylogenetic tree of Pol protein sequences (n = 111; see Supplementary Table S1) was constructed (1,000 bootstrap replicates). Annotations of viral strains and GenBank IDs are colored according to the types: *vpu*⁻*vpx*⁻ (black), Vpx-type (orange), Vpu-type 1 (red), Vpu-type 2 (brown), Vpu-type 3 (green), and Vpu-type 4 (blue). Both HIV-1 (Group M) and HIV-2 were collapsed and displayed as black triangles. The scale bar below the tree indicates 0.09 (9%) amino acid substitutions per site, and bootstrap values (1/100), indicating estimated posterior probabilities, are given at each node.

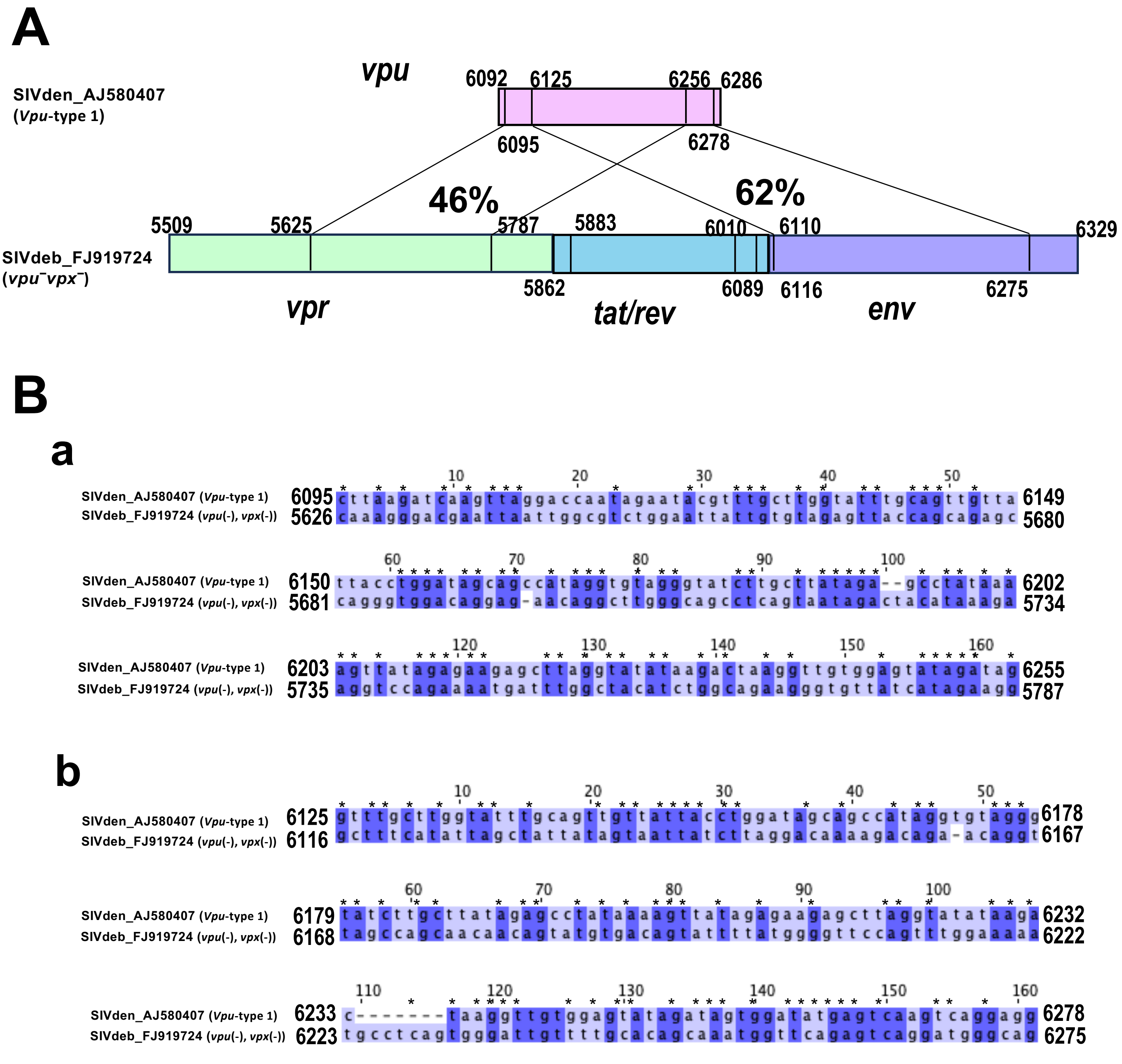


**Supplementary Figure S5.** Sequence similarities between SIVden *vpu* and SIVdeb *vpr* and *env: Case 1*. (A) Schematic representation of the viral gene structures of *vpu*, *tat-rev*, *vpr*, and *env*. The exact position of each gene within the genome is indicated. The vertical lines within the boxes represent the locations of homology between the genes. (B) Nucleotide sequence alignments of (a) *vpu* and *vpr*, and (b) *vpu* and *env*. '*' indicates nucleotide conservation.

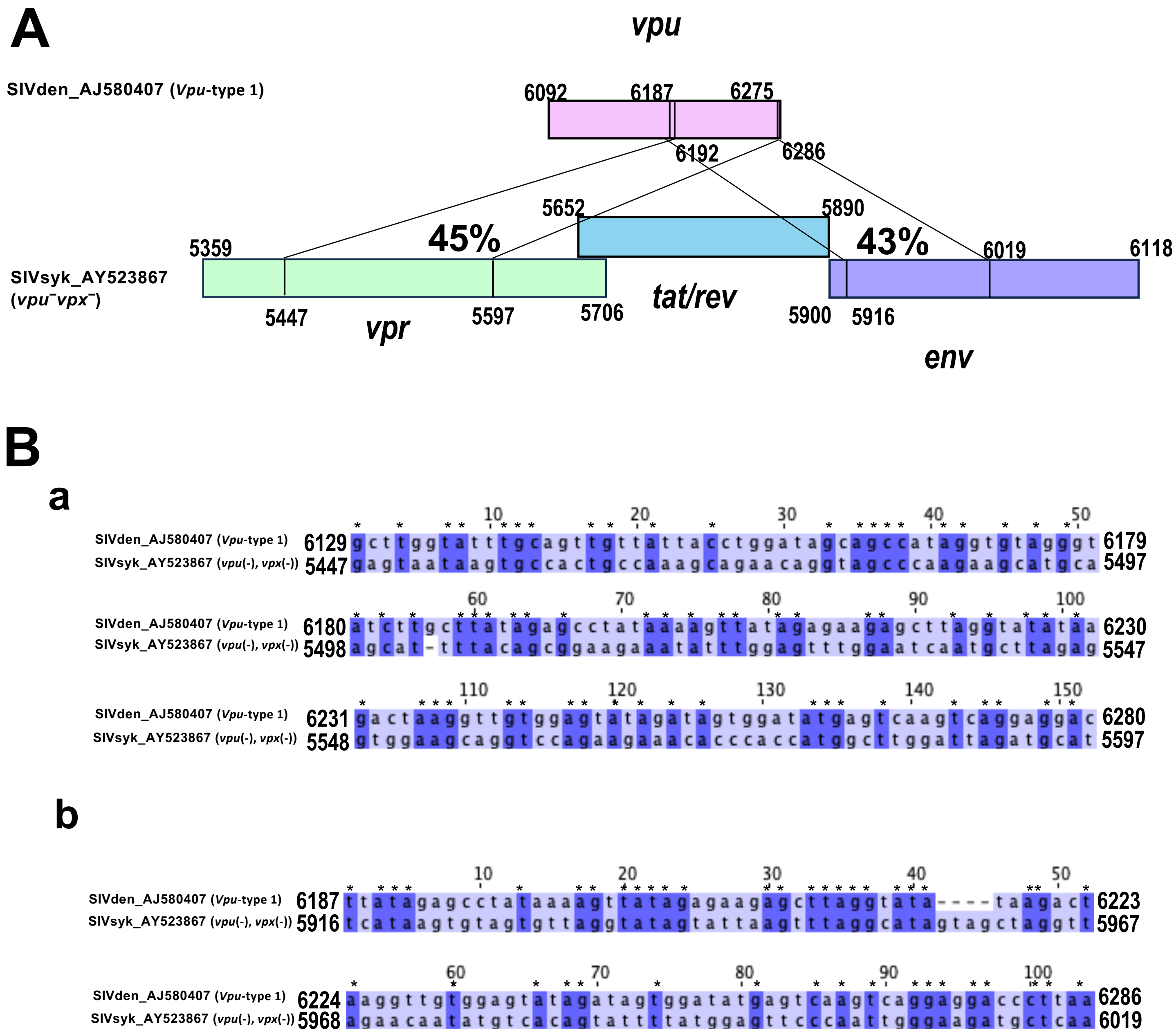


**Supplementary Figure S6. Sequence similarities between SIVden *vpu* and SIVsyk *vpr* and *env: Case 2.*** (A) Schematic representation of the viral gene structures of *vpu*, *tat-rev*, *vpr*, and *env*. The exact position of each gene within the genome is indicated. The vertical lines within the boxes represent the locations of homology between the genes. (B) Nucleotide sequence alignments of (a) *vpu* and *vpr*, and (b) *vpu* and *env*. '*' indicates nucleotide conservation.

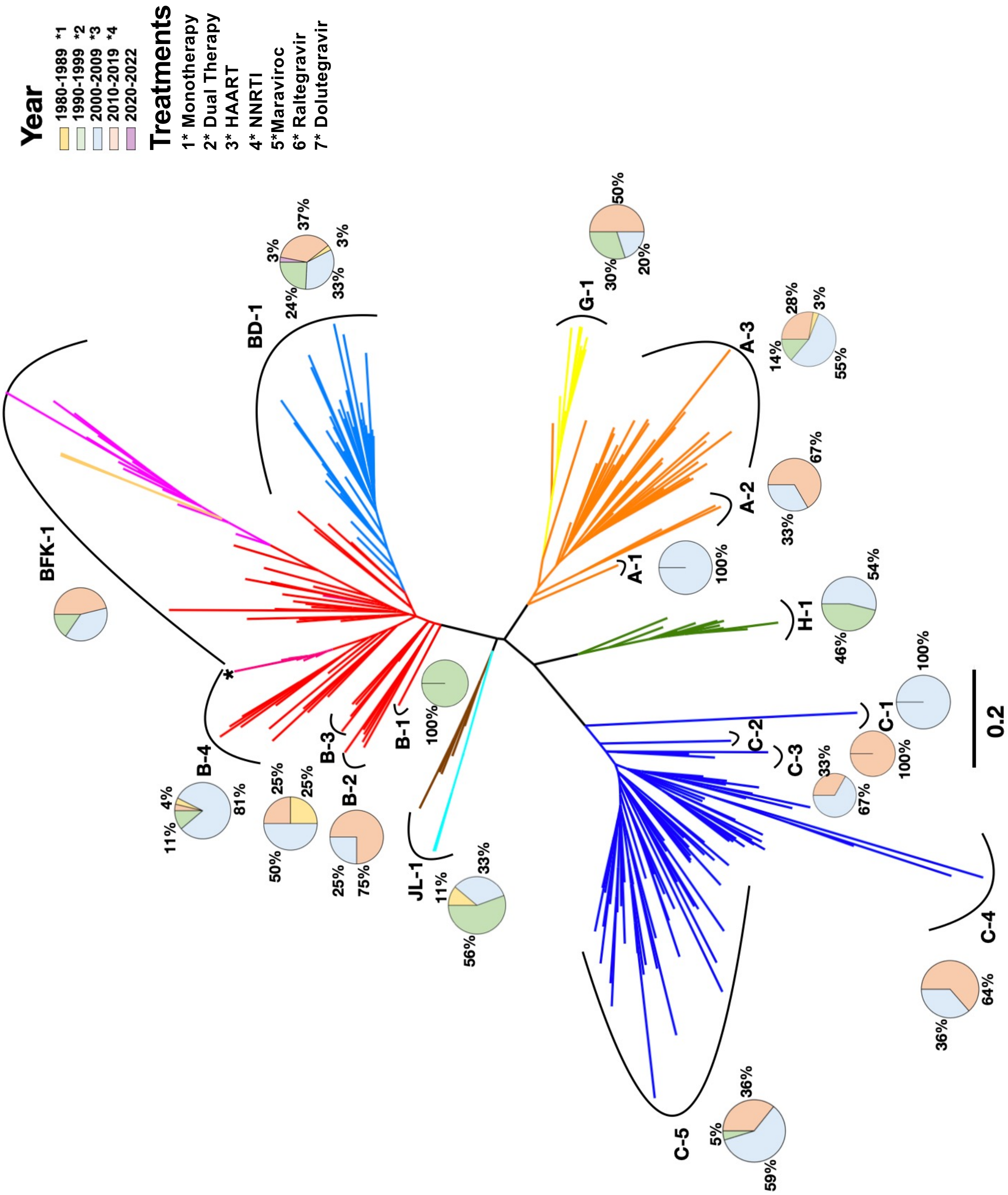


**Supplementary Figure S7.** Unrooted phylogenetic tree of Vpu proteins and the temporal (decadal) distribution of HIV-1 group M Vpu. The unrooted phylogenetic tree was constructed with Vpu protein sequences (n = 242; see Supplementary Table S1) (1,000 bootstrap replicates). The branches of the phylogenetic tree are colored according to their official viral subtype. The pie chart shows the temporal prevalence of the protein subtypes. Years are colored on a 10-year basis: 1980–89 (yellow), 1990–99 (green), 2000–09 (blue), 2010–19 (orange), and 2020–22 (purple). '*' indicates when each treatment was approved by the US Food and Drug Administration. 1* zidovudine (1987), 2* saquinavir (1995), highly active anti-retroviral therapy (HAART) (1996), non-nucleoside reverse transcriptase inhibitors (NNRTI) (1996), lamivudine/zidovudine (1997), 3* integrase inhibitor raltegravir (2007), CCR5-blocking drug maraviroc (2007), 4* pre-exposure prophylaxis (PrEP) (2012), and second-generation integrase inhibitor dolutegravir (2013). The scale bar below the tree indicates 0.2 (20%) amino acid substitutions per site.

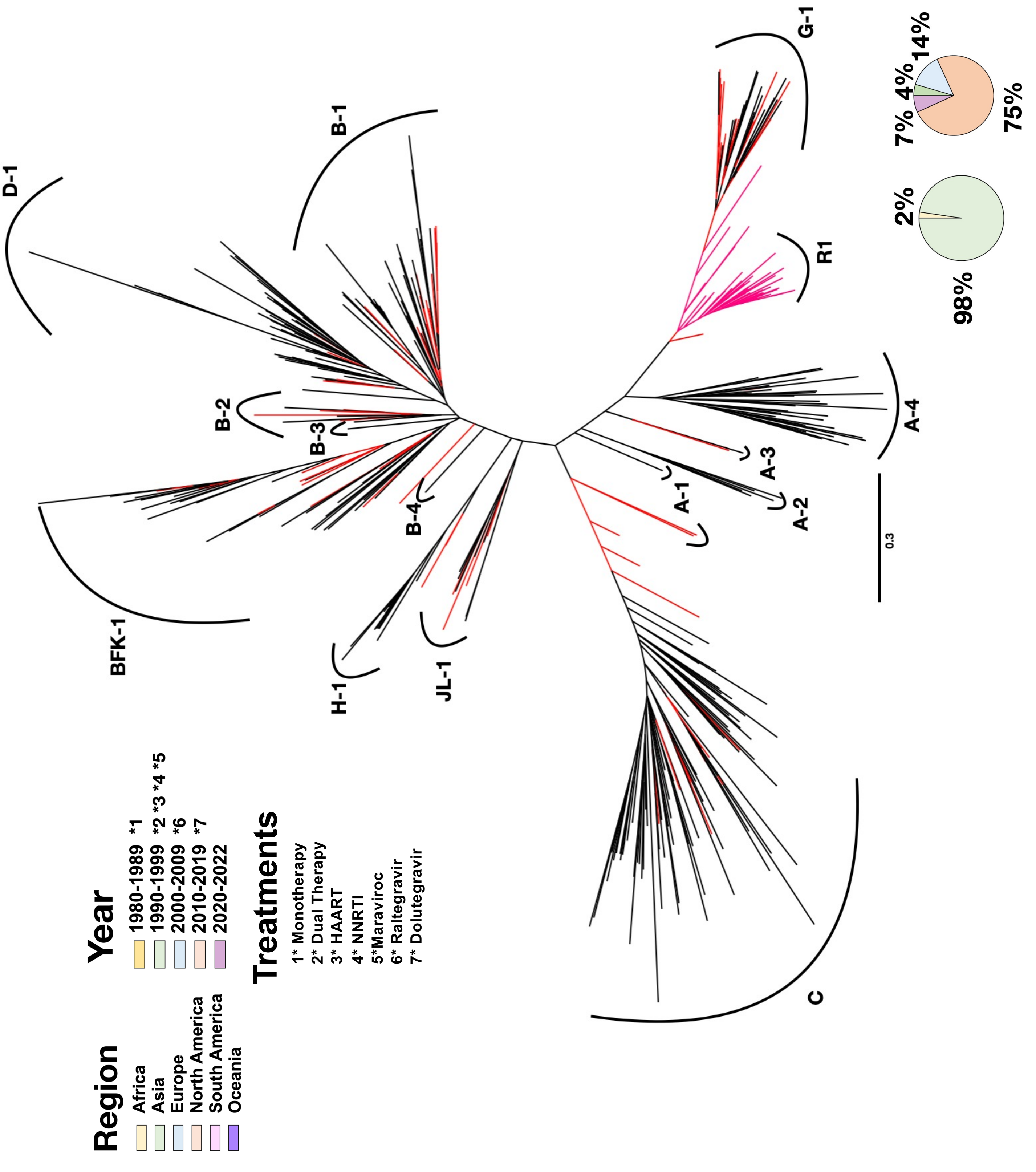


**Supplementary Figure S8.** Unrooted phylogenetic tree of the circulating recombinant forms of Vpu proteins. The unrooted rooted phylogenetic tree of Vpu protein was constructed, incorporating sequences from Supplementary Fig. S7 together with an additional set of 135 circulating recombinant form sequences (total dataset, n = 387 sequences). R1 is a classified clade containing only recombinant forms. All the circulating recombinant forms are colored red, and the R1 cluster is colored pink; other viral subtypes are colored black. Pie charts of the regional (left) and temporal (decadal) distributions (right) of R1 are located in the bottom right corner.

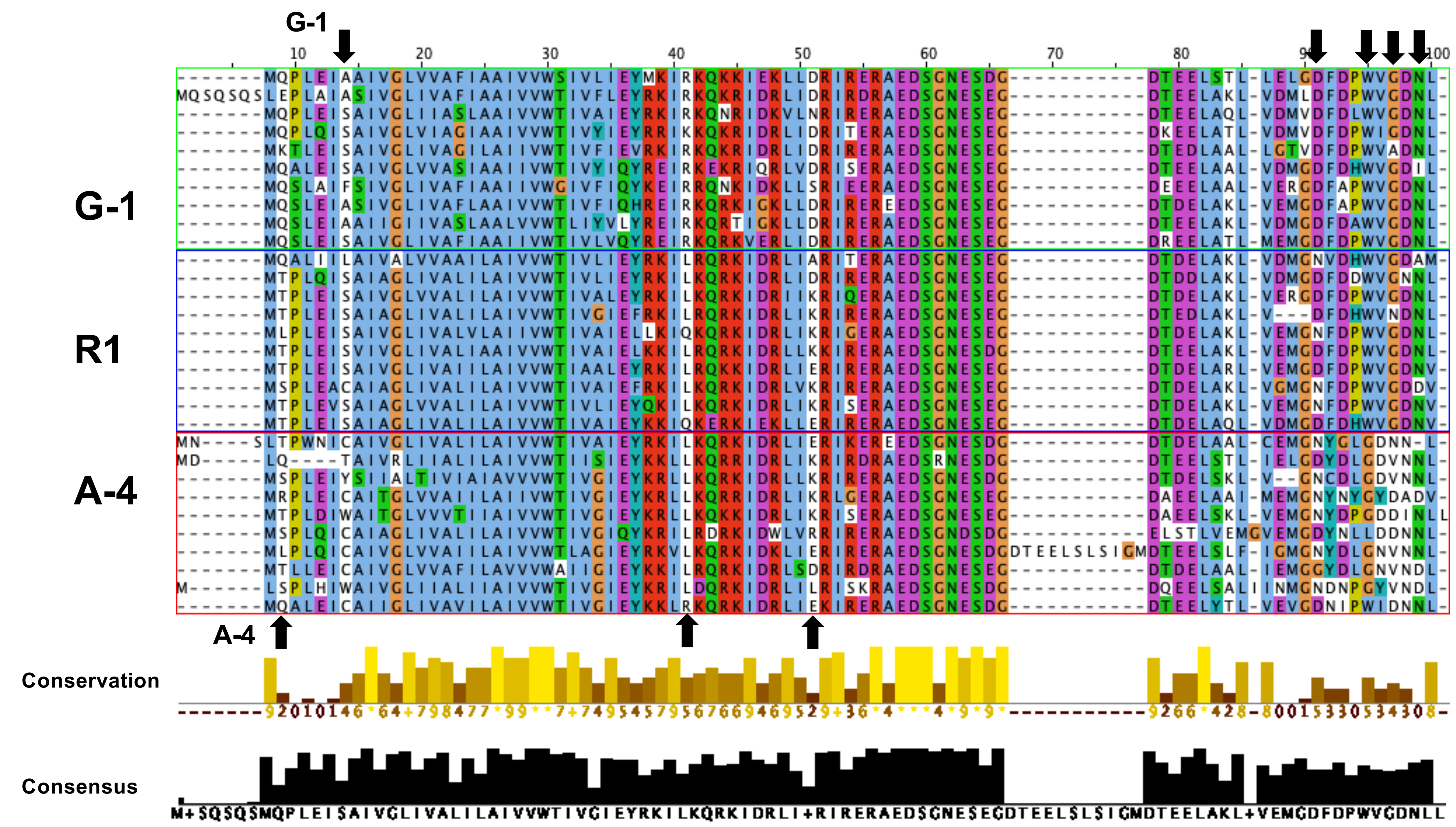


**Supplementary Figure S9.** Multiple-amino-acid-sequence alignment of Vpu proteins of G-1, R1, and A-4. A multiple-sequence alignment of each type of Vpu protein, colored according to the ClustalX color scheme (n = 30). Histogram below the alignment describes the sequence conservation of the residues according to their physiochemical properties. The consensus sequence shows conserved regions. The nonconserved residues are marked with '+'. The upward arrows indicate residues that R1 shares with A-4 and downwards arrow indicate residues it shares with G-1.

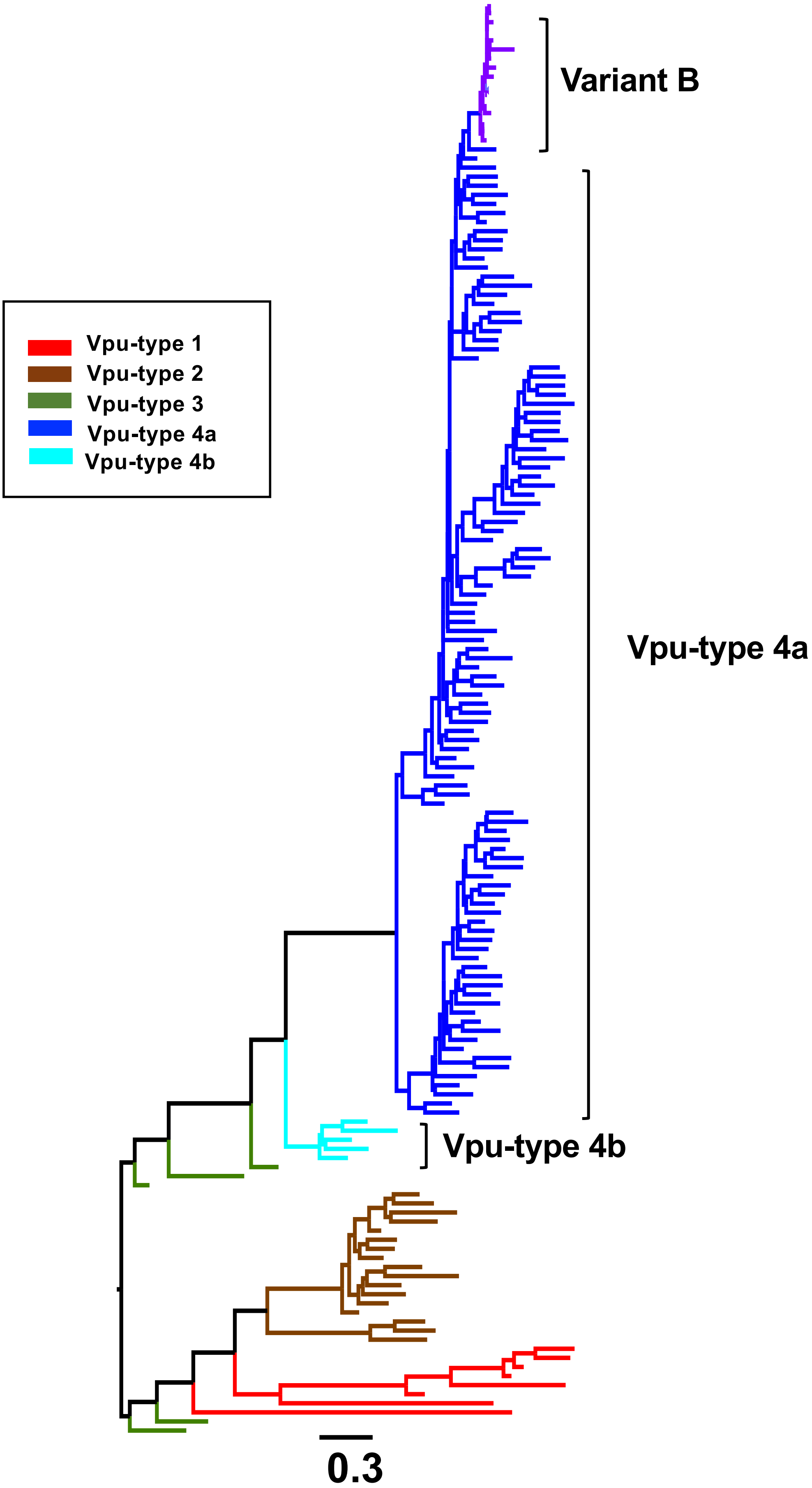


**Supplementary Figure S10.** Rooted phylogenetic of Vpu protein sequences of uncollapsed HIV-1 and SIV strains. The midpoint-rooted phylogenetic tree of Vpu proteins was constructed (1,000 bootstrap replicates) by incorporating sequences from Fig. 1, together with an additional set of 17 variant B sequences (total dataset, n = 158 sequences). The branches are colored according to the virus type: Vpu-type 1 (red), Vpu-type 2 (brown), Vpu-type 3 (green), Vpu-type 4a (bright blue), and Vpu-type 4b (cyan). Variant strain B is colored purple. The scale bar below the tree indicates 0.3 (30%) amino acid substitutions per site, and bootstrap values (1/100), indicating estimated posterior probabilities, are given at each node.

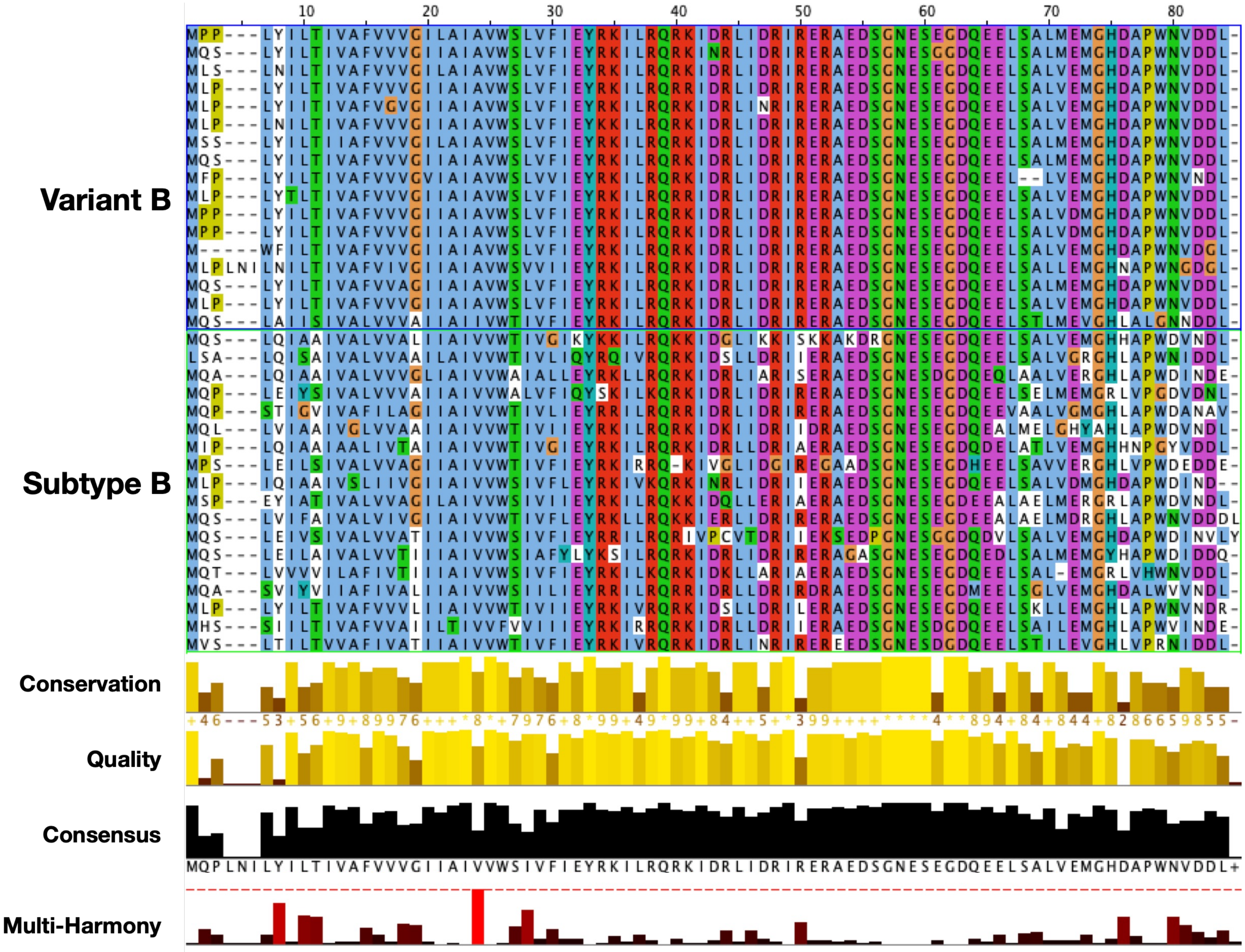


**Supplementary Figure S11.** Specificity analysis of variant B and subtype B. Multiple-sequence alignment of subtype B and variant B strains colored according to the ClustalX color scheme. The histogram below the alignment describes the conservation of residues according to their physiochemical properties, the quality annotation represents the likelihood of observing a mutation in any particular column of the alignment based on BLOSUM62 matrix scores. The consensus sequence shows the conserved regions. The nonconserved residues are marked with '+'. The multi-Harmony histogram shows subtype-specific residues.
